# High-density extracellular recordings from the interior of intact brain organoids enable automated high-throughput functional assay

**DOI:** 10.1101/2025.09.12.675888

**Authors:** Fan Wu, Cuthbert Steadman, Liam Argent, Brian Jamieson, Shamsudheen Karuthedath Vellarikkal, Alex M. Tamburino, Iya Khalil, Vanessa M. Peterson, Matthew E. Kennedy, Rebecca Mathew, Eran Stark, Bilal Cakir

**Author notes:** Correspondence (R.M.), (E.S.), (B.C.).

## Abstract

Bridging the gap between preclinical screening and clinical outcomes remains a major challenge in drug development for neurological disorders. Brain organoids, derived from human induced pluripotent stem cells, offer a scalable and physiologically relevant platform to model human neural circuits. We develop a fully automated system to record neural activity from the interior of intact human cortical organoids using a high-density microfabricated probe. The robotic system completes insertion within minutes while preserving organoid integrity and enables immediate recording of spontaneous spikes. We extract physiologically grounded and deterministic spike features, and train a long short-term memory classifier to distinguish between organoids derived from healthy individuals and those harboring familial Alzheimer’s disease (AD) mutations in the amyloid precursor protein. Despite intra-class variability, the classifier differentiates between organoid classes with 100% accuracy. The model classifies AD organoids treated with a drug candidate that reduces amyloid-β levels as retaining an AD-like electrophysiological phenotype, demonstrating that functional readout can contradict molecular markers. The findings establish a high-throughput functional framework that may complement and extend existing drug screening assays.

## INTRODUCTION

Improving the translation of preclinical screening assays into clinical outcomes remains a central challenge in the development of therapies for neurological diseases^1–3^. Recently, in vivo studies have emphasized the importance of circuit-level phenomena in neurological diseases^4–6^. However, traditional animal electrophysiology studies often require time-intensive surgeries and analyses, are difficult to scale, and are unsuitable for drug screening, where automation and high throughput are critical. Effective tools to bridge this gap are lacking.

Brain organoids derived from human induced pluripotent stem cells (hiPSCs) have emerged as a promising in vitro model, offering scalability, genetic fidelity, and the potential to recapitulate human-specific neurodevelopmental and disease-relevant features^7–9^. Unlike animal models, organoid models are versatile, potentially enabling large-scale drug screening that is not feasible with labor-intensive animal studies. Moreover, physiological data from human-based organoids can be more translatable than those from animal models, which often fail to predict human responses^10,11^. Organoids are three-dimensional (3D) structures that mimic diverse brain region identities, establish synaptic networks, and achieve increasingly mature phenotypes^12–17^. Despite these advantages, the functional interrogation of brain organoids remains underdeveloped, particularly with respect to electrophysiology.

Presently, functional electrophysiology in brain organoids primarily utilizes patch clamp and surface microelectrode arrays (2D-MEAs)^18–21^. Single electrode patch clamp techniques offer focal intracellular readout from individual cells, but lack scalability in terms of simultaneously recorded cells, limiting their utility for circuit or network-level analysis. While initially restricted to academic usage due to their high complexity and low throughput, patch clamp techniques saw widespread industrial adoption following the development of automated technologies^22,23^. By contrast, 2D-MEAs can include thousands of electrodes^24–27^, enable detection of synchronous neuronal firing across a large surface area^19,28^, and show spontaneous single unit firing sequences during neurodevelopment^29^. However, 2D-MEAs are by definition planar and optimized for 2D cultures^30^. When applied to 3D cultures, 2D-MEAs primarily sample from neurons near the tissue surface or outgrown neurites, providing limited access to deeper neuronal populations^31,32^. The employment of 2D-MEAs for recording from the depth of organoids often requires slicing or dissociation procedures that compromise tissue integrity^20,33^.

The limitations of 2D-MEAs for 3D cultures were recognized early in brain slice electrophysiology^34,35^, motivating the development of microfabricated MEAs with protruding or 3D electrode geometries designed to improve cell–electrode coupling through superficial dead-cell layers^34,35^. Although these approaches enhance signal quality near the tissue surface, they remain constrained by fixed electrode heights which limit penetration depths^21^. Ongoing efforts aim to increase electrode length^36^ and introduce tunability^37^. However, such advances substantially increase microfabrication complexity, reduce manufacturing yield, and raise cost, thereby limiting scalability and hindering broad commercialization.

These challenges are further amplified in brain organoids, which are substantially thicker than acute slices and continue to increase in size and structural complexity. Advances such as vascularization strategies aim to support larger organoids by mitigating diffusion-limited nutrient and waste transport from the interior^38^. As organoid technology evolves and organoid sizes expand, effective functional interrogation increasingly requires electrophysiological access to deep, interior neuronal populations without compromising tissue viability or architecture. To fully leverage the 3D cytoarchitecture of brain organoids, functional recordings must access their interior non-destructively. Two general strategies have been developed to address this challenge. The first approach employs flexible or mesh-like electrode arrays introduced at early stages of organoid formation, allowing neurons to grow into three-dimensional structures around the electrodes over time^39,40^. This strategy enables minimally invasive, longitudinal recordings across organoid development and maturation.

The second approach for functional interrogation of organoids involves microfabricated silicon probes. Silicon probes are rigid depth electrodes that can be inserted into mature structures to record neural activity at user-defined depths^41,42^. A single probe can be used repeatedly to sequentially sample multiple organoids, making silicon probes well suited for acute recordings and high-throughput experimental paradigms. Silicon probes microfabricated by advanced semiconductor processes have long been used in systems neuroscience to record from live animals, offering dense, minimally invasive, multichannel access to circuit-level dynamics in deep brain structures^41,42^. While multi-channel silicon probes are indispensable in neuroscience research, their adoption in industrial and translational settings remains limited due to the manual, low-throughput nature of surgeries and the technical burdens of analysis. Several recent studies have successfully demonstrated the feasibility of using silicon probes in organoids^19,43–45^.

Here, inspired by the analytical power of silicon probes for depth recordings and the transformative impact of automation for patch clamp techniques, we developed a fully automated platform for high-throughput, depth-resolved electrophysiology in intact brain organoids. Our system automates the entire pipeline: from micrometer-precision probe insertion, through spontaneous multi-site spike recording, and all the way to feature extraction and machine learning-based classification at the system level. We demonstrate the utility of this approach by recording from human cortical organoids (hCO) with and without known genetic perturbations, namely isogenic Alzheimer’s disease (AD) model carrying the amyloid precursor protein (APP) Swedish (KM0/671NL) and Indiana (V717F) mutations^46^. Despite inherent biological variability, electrophysiological features vary more between organoid classes than within each class. Machine learning classifiers accurately predict organoid class identity from short novel recordings. These results establish the feasibility of a rapid functional assay, providing a framework for integrating electrophysiology-based phenotyping into high-throughput preclinical pipelines.

## RESULTS

### An automated instrument allows recording from the inside of brain organoids

To enable high-throughput and consistent recordings from within intact brain organoids, we developed a fully automated instrument capable of precisely inserting a high-density neural probe into the organoid interior and acquiring spontaneous electrophysiological activity. The system is housed in a custom chamber that provides electromagnetic shielding, with internal modules for organoid confinement, probe actuation, and recording (**Fig. 1A**).

**Figure 1.**
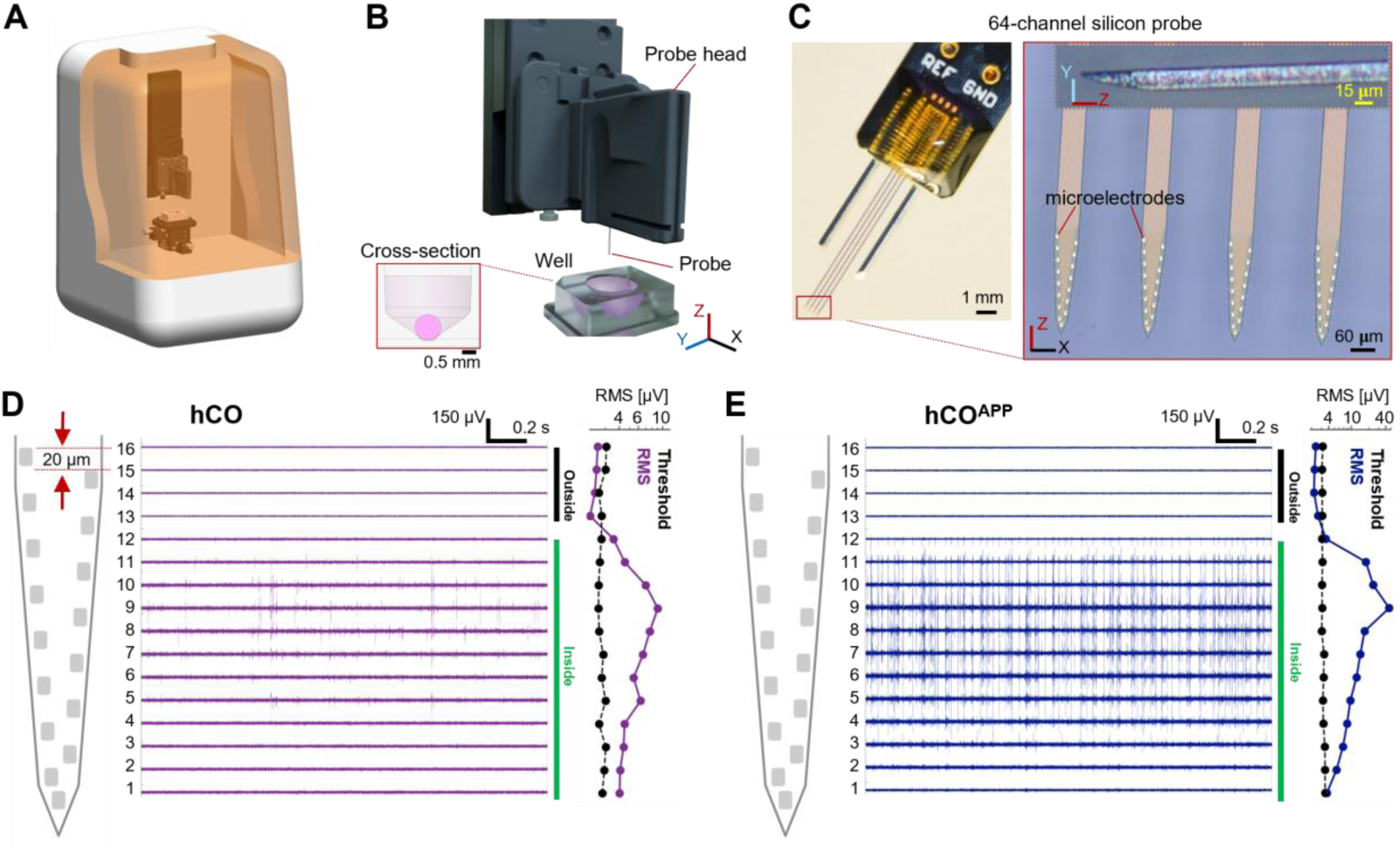
High-throughput instrument automatically inserts a 64-channel microfabricated neural probe and immediately records spiking activity from the inside of intact organoids. (A) External design of the instrument chamber. (B) Internal design, showing a drivable probe head and a custom designed stationary well. (C) Left, Four-shank microfabricated probe assembled on a printed circuit board. Right, Magnified view of the probe tips showing 64 gold microelectrodes integrated on four penetrating probe shanks with sub-micrometer precision. Inset, Sharpened shank tip. (D) Recording from a representative human cortical organoid (hCO). Spikes are recorded immediately after insertion and are concentrated on a continuous subset of recording electrodes that are inside the intact organoid. Left, Shank schematic showing 20 μm spacing along the Z-axis between adjacent electrodes. Middle, Bandpass filtered (300-6,000 Hz) traces across one probe shank over a 2 s time window. Right, V_RMS_ (purple) and pre-computed thresholds (black) for every channel distinguish electrodes inside from outside of the organoid. (E) Same as D, for a representative hCO mutant for the amyloid precursor protein (hCO^APP^).

The core of the system consists of a custom-designed well and a motorized probe head (**Fig. 1B**). The well is 9 mm deep, featuring conical geometry with a 45° taper and a 0.5 mm-diameter base, engineered to capture organoids with diameters ranging 0.5-2 mm. This geometry ensures that organoids dropped from a pipette anywhere within the well will fall to its center by gravity. The slanted walls provide lateral support that confines and stabilizes the organoid during probe insertion and recording. Inside the enclosure, the probe head incorporates a microfabricated silicon neural probe (P64-1, Diagnostic Biochips, Glen Burnie, MD) and an integrated amplifier (RHD2164, Intan Technologies, Los Angeles, CA), mounted on a precision motorized single-axis actuator (KMTS50E, Thorlabs, Newton, NJ). After calibration, the assembled mechanical system maintains ≤100 µm lateral misalignment in the X/Y directions, 0.8 µm Z direction steps, and 15 µm repeatability in the Z direction.

The silicon probe comprises four 9 mm-long shanks, each carrying 16 microelectrodes (**Fig. 1C, left**). Recording sites distribute across a 750 µm lateral span (X axis) over the four shanks, and over 300 µm depth (Z axis) per shank (**Fig. 1C, right**). This coverage is well-suited to the macroscopic organization of many cortical organoids, which are approximately spherical with diameters of 0.5–2 mm. Although early-stage organoids contain multiple radial rosettes from which neurons migrate outward, these structures become less distinct as organoids mature, and differentiated neurons tend to form a more continuous, neuron-rich outer layer (**Fig. S1A**). At the same time, multiple studies have shown that reduced activity and cell loss toward the core primarily reflect hypoxia, stress, and necrosis rather than an absence of neurons, with characteristic hypoxic length scales on the order of a few hundred micrometers in conventionally grown whole organoids^19,47–51^. The multi-shank probe used here spans ∼300 μm along the Z axis, which enables placing most recording sites within the outer neuronal shell while avoiding the deeper, low-activity core (**Fig. 1D-E**). Standard silicon shanks with a uniform cross-section (roughly 60 µm-wide, XZ plane; and 15 µm-thick, YZ plane) are routinely used to penetrate the rodent pia mater^52^. While this form factor allowed penetrating organoids, we found it essential to sharpen the tips along the YZ plane to minimize tissue compression just before the probe tips penetrate (**Fig. 1C right, inset**). The sharpened tips improve recording stability and allow reliable control of probe descent with predictable signal changes in response to incremental depth adjustments. Additionally, post-recording staining shows intact, healthy neuronal shell (**Fig. S1B**). The median caspase-3/DAPI ratio in 150-day-old organoids post 1-hour recordings is 30 (IQR, [29.0 31.5]; n=16), not significantly different from pre-recording 150-day-old organoids (median: 30.0; IQR, [28.5 31.0]; n=16; p=0.74, Mann-Whitney U-test), indicating that the microfabricated silicon shanks do not measurably compromise the acute health of neuronal tissue in these cortical organoids.

### V_RMS_-based algorithm enables closed-loop and precise probe insertion into organoids

To optimize electrode placement within active neuronal regions of intact brain organoids, we developed an automated insertion algorithm guided by real-time measurements of the root mean square (RMS) extracellular voltage (V_RMS_). During the initial phase of insertion, the Z axis manipulator lowers the probe to a depth above the organoid, until all electrodes are fully immersed in media (BrainPhys, STEMCELL Technologies, BC, Canada). At this stage, we establish a baseline V_RMS_ for each recording channel by computing the mean and standard deviation (SD) of 100 consecutive V_RMS_ samples (one sample every 100 ms) from band-pass filtered (300-6,000 Hz) signal from that channel. Then, a media-organoid boundary detection threshold is defined for each channel as five SDs above the mean.

In the second phase, the Z axis manipulator advances the probe in 50 µm steps, pausing for 3 seconds at each step to acquire a new set of 30 V_RMS_ measurements. As the probe enters the organoid, the bottom-most electrodes begin transitioning from the media into the tissue and reliably exhibit a characteristic increase in V_RMS_. Once this transition is detected at the bottom-most electrodes, the step size is reduced to 20 µm, the vertical spacing between adjacent electrodes. An electrode is marked “inside” when its updated V_RMS_ exceeds its own out-of-organoid pre-computed threshold. This incremental, depth-resolved approach enables the algorithm to track the progressive recruitment of electrodes and to halt insertion once the majority of the electrodes are inside the organoid, indicating optimal probe placement.

The elevated V_RMS_ inside the organoid does not require the presence of local spiking activity; it likely reflects the presence of neurons and neuronal processes (axons and dendrites) and their subthreshold signals (**Fig. 1D-E; Fig. S2A**). The higher impedance of the tissue compared with the fluid medium contributes to the effective impedance of the electrodes, increasing the Johnson noise (1/f noise component is negligible in the 300- 6,000 Hz range), further increasing the RMS. To isolate these effects from spike contributions, we excluded spike samples (40 samples centered around every spike extremum) and then computed an RMS value within the first 0.1 s window. Log-transformed RMS values from 6,656 electrodes across 120 recordings were fit with a two-component 1D Gaussian Mixture Model (**Fig. S2B**). Bayesian Information Criterion for one-, two- and three component models are 27749.38, 18818.15, and 17720.20, respectively, with substantial improvement when increasing the model from one to two Gaussian components, supporting the presence of at least two separable RMS populations. One Gaussian (mean=0.366; SD=0.042; n=4,073; weight=0.591) approximates log-RMS distribution of electrodes in the media (outside organoid), while the other Gaussian (mean=0.677; SD=0.157; n=2,583; weight=0.409) reflects electrodes inside organoids (**Fig. S2B**). The clear separation between the two Gaussians demonstrates that V_RMS_ can reliably indicate electrode penetration into an organoid even in the absence of spontaneous spiking.

To illustrate the RMS changes in depth including contribution by spikes, **Figure 1D** shows the probe shank schematic (left), with each electrode aligned to the corresponding voltage time series (middle) and V_RMS_ measurements (right). In this representative recording from an hCO, the four top channels (13-16) remain in the media, while the other 12 channels penetrate the organoid, as indicated by their V_RMS_ values exceeding the predefined thresholds. Notably, channel 12 marks the uppermost electrode inside the organoid, but is too distant from active neurons to clearly detect spikes. Channels 5-11 display the most robust spiking activity, corresponding to a further increase in V_RMS_. Channels 1-4 are deepest within the organoid, showing sufficiently high V_RMS_ to be labeled as “inside”, but do not monitor spikes. When electrodes are advanced beyond this depth, V_RMS_ occasionally reduces to near-media levels (not shown). These findings suggest that the most reliable method for targeting active neurons is to maximize the number of electrodes positioned within the outer neuronal shell of the organoids (150–300 µm), using V_RMS_ feedback to avoid overshooting and passing this layer.

As a second example, **Figure 1E** illustrates a recording from an organoid of the same cell line that has been modified to incorporate APP mutations (hCO^APP^). In the hCO^APP^ preparation, rapid and spatially dense spiking was observed. The activity was concentrated between channels 2-12, in stark contrast to the relatively less active band observed in the hCO (**Fig. 1D**). Statistical analysis of the spike features from multiple hCOs and hCO^APP^s are described later in **Table 1** and **Fig. 5** (n=89 sessions; each session is a recording from one unique organoid). However, in both preparations, the thickness of the electro-physiologically identified neuronal shell was similar, corresponding to about 200 µm. In both preparations, the RMS-based algorithm was effective in reliably positioning the probe within the active layer, characterized by clear signal boundaries above and below.

**Table 1.** Feature statistics for every organoid class. ^a^Values indicate median [IQR] over all spikes.

| Organoid class | Sessions | Spikes | Amplitude<br>[μV]<sup>a</sup> | Peak A<br>[μV] | ISI<br>[ms] | TPT<br>[ms] |
| --- | --- | --- | --- | --- | --- | --- |
| hCO | 24 | 2,282,223 | -58.78<br>[36.64] | 8.91<br>[15.99] | 174<br>[304.5] | 0.48<br>[0.3] |
| hCO<sup>APP</sup> | 33 | 10,464,522 | -76.7<br>[69.72] | 11.94<br>[25.72] | 42.1<br>[82.55] | 0.38<br>[0.28] |
| hCO<sup>APP</sup> +<br>10 nM MK-8722 | 12 | 6,516,509 | -63.06<br>[41.85] | 12<br>[17.01] | 58.35<br>[162.05] | 0.43<br>[0.26] |
| hCO<sup>APP</sup> +<br>100 nM MK87-22 | 13 | 6,099,906 | -67.6<br>[53.62] | 14<br>[26.04] | 38.55<br>[56.6] | 0.39<br>[0.36] |
| hCOL<sup>APP</sup> | 7 | 452,151 | -65.1<br>[32.02] | 8.84<br>[14] | 51.1<br>[204.7] | 0.43<br>[0.3] |
| Summary | 89 | 25,815,311 | -68.5<br>[53.53] | 11.97<br>[21.97] | 46.8<br>[115.1] | 0.39<br>[0.3] |

### hCO variants permit disease modeling and assessment of drug treatments

We generated hCOs using an established protocol^53^, and selected distinct variants to test the sensitivity of our electrophysiology platform around day 150 (**Fig. 2A**). Consistent with prior studies^7,48,49^, immunofluorescence analysis showed strong expression of the excitatory neuron marker vesicular glutamate transporter 2 (VGLUT2), while the inhibitory neuron marker vesicular GABA transporter (VGAT) remained limited on day 120, consistent with a predominantly excitatory neuronal identity (**Fig. S3A**).

**Figure 2.**
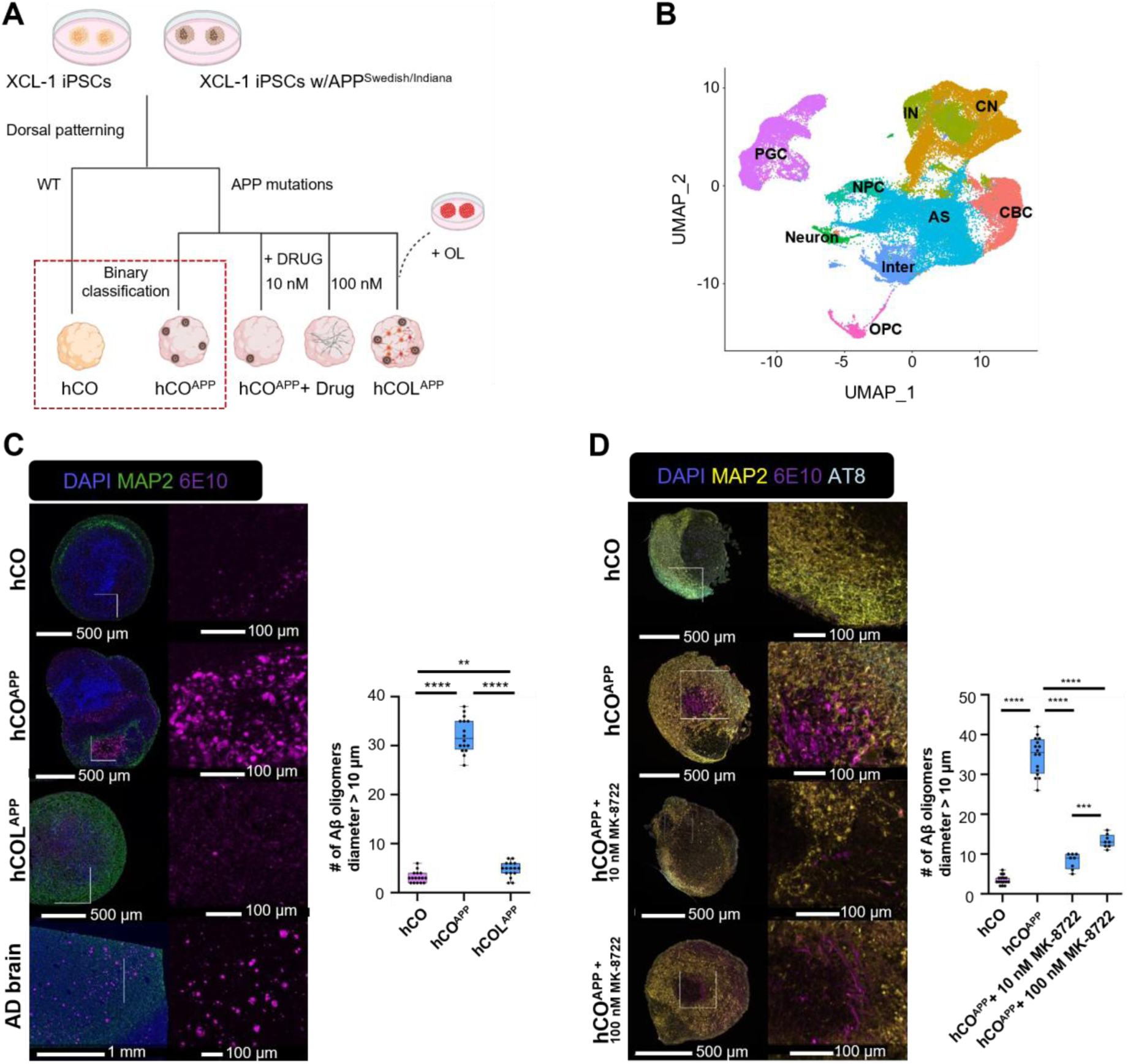
Generation and characterization of human cortical organoid and APP mutants. (**A**) Schematic representation of the generation of isogenic variants of hCOs and the treatments used in this study. hCO and hCO^APP^ are used for the binary classification task. OL, oligodendrocytes. (**B**) UMAP projection of single-cell RNA sampled at day 120 reveals nine distinct clusters, corresponding to distinct cell types, including neural progenitor cells (NPCs), cortical neurons (CNs), interneurons (INs), astrocytes (AS), oligodendrocyte progenitors (OPC), intermediate cells (Inter), proteoglycan expressing cells (PGC), and cilia-bearing cells (CBC). (**C**) Left, Immunostaining of hCO and hCO^APP^ variants at day 120, and AD patient post-mortem brain for neuronal marker MAP and Aβ antibody 6E10. Right, Quantification of Aβ aggregates in organoid variants. (**D**) Left, Co-staining of Aβ (6E10), phospho-tau (AT8) and neuronal marker MAP2 in hCO and hCO^APP^ with and without drug treatments (10 nM and 100 nM MK-8722) at day 150. Right, Quantification of Aβ aggregates in organoid variants. In **C** and **D**, box plots show medians and interquartile range over three independent batches, n=16 organoids in each group. **/***/****, p<0.01/0.001/0.0001, two-tailed Mann-Whitney U-test.

To assess cellular diversity and the maturation process, we performed single-cell RNA sequencing at days 50 (n=10) and 115 (n=6; **Table S1**). Dimensionality reduction using uniform manifold approximation and projection (UMAP) revealed nine transcriptionally distinct clusters, which we classified as neural progenitor cells (NPCs), cortical neurons (CNs), interneurons (INs), astrocytes (AS), and five other previously described^38,54^ populations (**Fig. 2B**; **Fig. S3B**, **S3C**). As a reference, we utilized the datasets from the Human Neural Organoid Cell Atlas (HNOCA) and its computational tools^55^, enabling us to automatically define the cell types present in the organoids and further validate their presence. The identity of the annotated cells was highly similar to the automated annotation prediction by the HNOCA model, suggesting that the cell types are indeed relevant to the forebrain **(Fig. S3D**). Thus, the hCO model recapitulates the cellular diversity observed in previously published cortical organoid models, providing an in vitro system for modeling human cortical development and function.

To test disease modeling using our recording platform, we developed hiPSCs carrying familial mutations in the APP (XCL-1_APP Ind (SwHomo)-3G05-E01) associated with early-onset AD^46^, and generated a baseline AD hCO (hCO^APP^). Given that deposition of amyloid beta (Aβ) and neurodegeneration occur sequentially in the human AD brain^56^, we examined Aβ accumulation in hCO and hCO^APP^ organoids at different developmental stages (**Fig. S4**). Compared with hCOs, the hCO^APP^ organoids exhibited excessive accumulation of Aβ aggregates (diameters above 10 µm) at all examined stages (days 50, 90, 120 and 150; p<0.001; n=16 hCO and n=16 hCO^APP^ analyzed at each developmental stage; two-tailed Mann-Whitney U-test; ***p<0.001), with the levels plateauing after day 120 (p=0.63, between hCO^APP^ at day 120 and day 150; **Fig. S4**). Consistent with the imaging data (**Fig. S4A-B**), ELISA measurements of formic-acid–extracted Aβ1-42 and Aβ1-40 (normalized to total protein) showed that hCO^APP^ organoids have a higher Aβ42/Aβ40 ratio in the insoluble fraction compared with control hCO samples (n=16 organoids per group; p<0.0001, two-tailed Mann–Whitney U-test; **Fig. S4C).** Notably, an increased Aβ42/Aβ40 ratio and robust Aβ aggregation have also been reported in familial AD human cerebral organoids, and our findings are consistent with these previously described AD-like pathological features^57–59^. To generate AD hCOs with oligodendrocytes (hCOL^APP^), we mixed 20% three-transcription factor (3-TF) induced control iPSCs with 80% APP mutation-containing iPSCs (**Fig. 2A**). At day 120, hCO^APP^ demonstrated a higher number of Aβ aggregates compared with hCO (n=16 hCO^APP^ and n=16 hCO; p<0.0001, two-tailed Mann-Whitney U-test; **Fig. 2C**, right). hCOL^APP^ (n=16) also showed elevated Aβ levels compared with hCO (p=0.004). Aβ levels in hCOL^APP^ remained lower than the levels in hCO^APP^ (p<0.0001), suggesting that the presence of oligodendrocytes may alter pathological progression.

To probe therapeutic responsiveness of hCO, we treated a subset of the hCO^APP^ with a beta-site APP cleaving enzyme 1 (BACE1) inhibitor, MK-8277^60^. We conducted electrophysiological recordings after 30 consecutive days of treatment using 10 nM and 100 nM concentrations. Immunofluorescence microscopy revealed a reduction in 6E10 antibody staining, confirming decreased Aβ accumulation in treated samples (p<0.001; n=16 organoids in each of the four groups; **Fig. 2D**). Because the hCO^APP^ line carries the APP Swedish mutation which has reduced sensitivity to BACE1 inhibition, these findings may be viewed as a pharmacological probe of assay responsiveness, rather than as a complete rescue of pathology. The results demonstrate that our disease-modeling platform recapitulates hallmark features of AD pathology and responds to pharmacological interventions.

### Depth-resolved spike-sorted units demonstrate recording quality and interior access in intact organoids

To validate recording quality and assess depth-resolved access in intact organoids, we performed spike sorting on all depth-probe recordings (n=89 organoids; **Table 1**) and analyzed the spatial distribution of the well-isolated single units identified from the high-density extracellular signals. Units were classified as single units based on an inter-spike interval (ISI) index below 0.2^61^. Accumulated over all 89 recordings, a total of 1,358 single units were identified.

For each unit, depth was defined relative to the media–organoid interface using the channel that exhibited the largest-amplitude waveform. Units were sorted by this peak-channel depth and summarized across recordings (**Fig. 3A**, left). For visualization, waveforms from each unit were collapsed into a single time-aligned representation, centered on the time of the waveform peak and displayed across channels along the probe shank.

**Figure 3.**
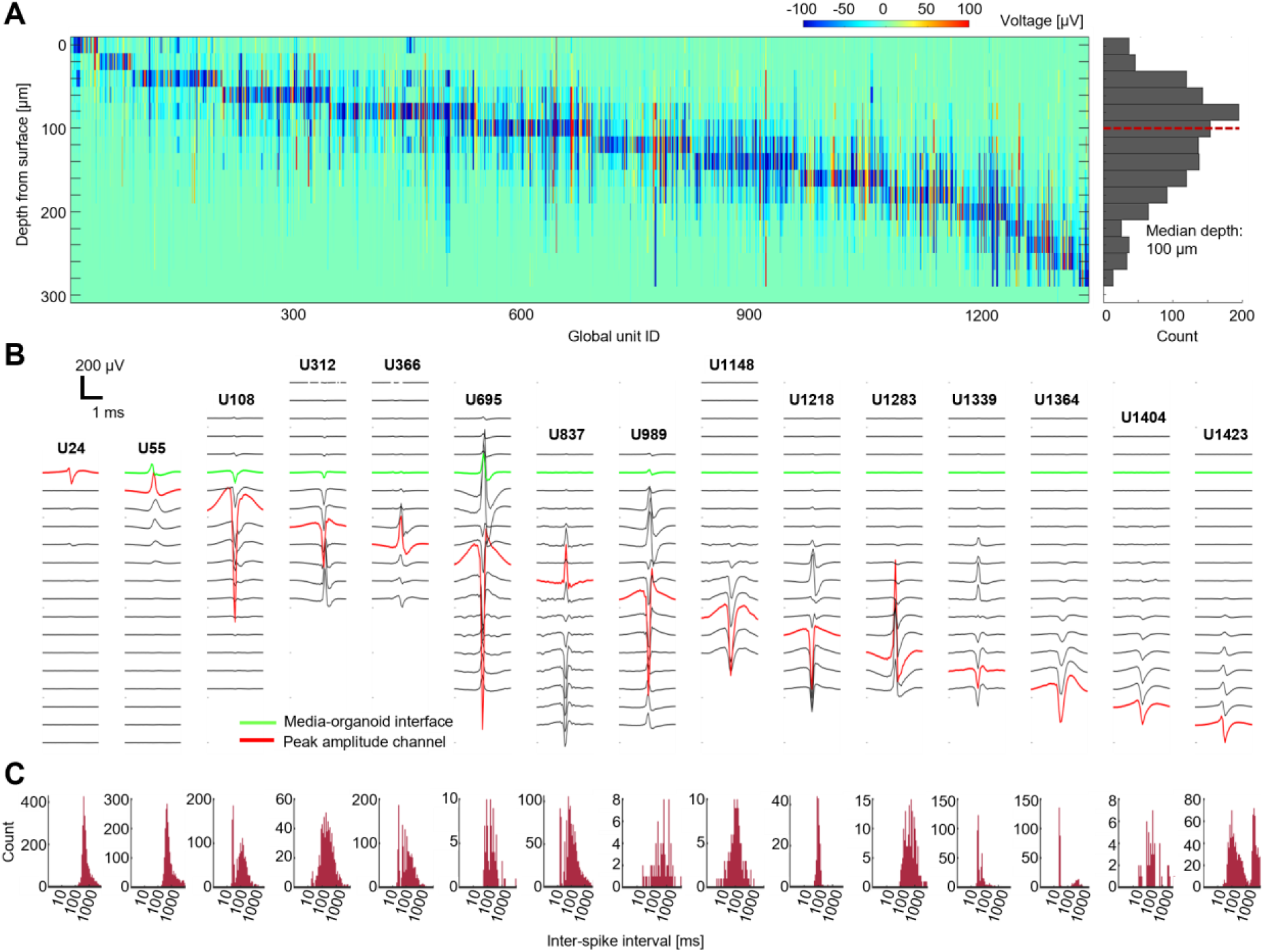
Depth-resolved single-unit recordings from high-density penetrating probes in intact brain organoids. (**A**) Left, Heatmap summarizing well-isolated single units (n=1,358) recorded from 89 organoids (**Table 1**), mapped to their relative depths from the media–organoid interface. Each column represents a single unit and displays the depth-resolved snapshot of its extracellular waveform, shown at the time point corresponding to the peak amplitude. Right, Histogram showing relative depths of the peak amplitude channels of all units. (**B**) Extracellular waveforms (band-pass filtered, 300-6000 Hz; 2.5 ms) from representative units spanning recording channels along the probe shank. Units are ordered from left to right by decreasing depth of the channel exhibiting the peak amplitude relative to the media–organoid interface. (**C**) Inter-spike interval histograms of the units shown in **B**.

The distribution of peak-channel depths shows a unimodal profile with a skew toward more superficial channels (**Fig. 3A**, right). The median depth of peak waveform channels was 100 µm, and the inter-quartile range (IQR) was [60 160] µm. The number of detected units increased from the surface to this median depth and decreased at greater depths. The decrease at deeper positions likely reflects probe insertion limits imposed by the automated stopping criteria and occasionally less active core (**Fig. 1DE**). To examine waveform properties across depth, we selected representative units with peak channels at discrete depths and displayed them (**Fig. 3B**). For each example, the peak channel is highlighted (red), along with the channel at the organoid surface (green) based on RMS threshold crossing. The depth difference between these two channels yields an estimate of the depth of the unit from the surface of the organoid (**Fig. 3A**). Both negative- and positive-polarity spikes corresponding to somatic spikes and return currents, respectively^62^, were observed. In some cases both polarities were detected within the same unit, corresponding to axonal spikes^62^ and highlighting the capability of the high-density electrodes to resolve subcellular structures. No systematic relationship between spike polarity and depth was observed, in contrast to the laminar polarity organization commonly reported in structured brain regions such as hippocampus or the neocortex.

ISI histograms for corresponding units in **Fig. 3B** show low refractory period violations, supporting the classification of these spike clusters as well-isolated single units. The ISI distributions reveal diversity in temporal firing patterns across units. Together, the features covered by waveform shape and spike timing demonstrate broad coverage of neurons recorded from hCOs, consistent with neuronal heterogeneity observed in cortical recordings from live subjects^63^, and highlighting the physiological relevance of intact brain organoids. These results suggest that depth-resolved extracellular recordings from intact organoids yield high-quality spike-derived features throughout the penetrated volume, providing a foundation for downstream analyses.

### Automated processing yields four deterministic spike features from raw recordings

An assay that enables high-throughput screening must extract meaningful patterns from neural recordings that distinguish control from test organoids accurately with minimal human intervention. Single-unit analyses (**Fig. 3**) can provide valuable insights into the identity, depth profile and dynamics of individual units and the neuronal circuitry. However, spike sorting and subsequent quality control, while increasingly supported by modern frameworks^64^, are still time-consuming, subjective, and require large numbers of spikes to reliably separate between spike clusters^65^. Moreover, stringent single-unit isolation criteria typically exclude a substantial fraction of detected spikes as contaminated multi-unit clusters. To enable scalable and reproducible functional phenotyping, we therefore implement a streamlined feature extraction pipeline that draws on insights from single-unit analyses without requiring any spike sorting.

We first bandpass-filter the raw data (300-6,000 Hz) on every channel and detect events using a 5 SD threshold from the mean. For each detected spike, we define the time of the spike as the maximum of the absolute value. In one representative hCO recording, 121 clusters (comprised of 453,995 spikes) are generated by Kilosort4^66^, of which 48 units (comprised of 202,393 spikes) are considered well-isolated single-units. In contrast, the threshold-based detection algorithm identifies 378,718 spikes, encompassing spikes corresponding to both well-isolated single-units and multi-units. When spike rasters of unsorted and sorted spikes from the four probe shanks are aligned, it is evident that in many cases, threshold-based spike detections are excluded when only well-isolated units are considered (**Fig. 4A**). Many spike events are detected on multiple channels at the same time, but for the purposes of feature extraction and session-level classification, they are not differentiated from isolated events (**Fig. 4B**).

1. For each unsorted spike, we extract a 2 ms waveform segment that starts 1 ms before the time of the spike (40 samples at 20 kHz). Then, we use the time of the spike and the extracted waveform to compute four features (**Fig. 4C, D**; **Table 1**). To provide an intuitive visualization of the feature space while maintaining consistency with the data structure used in the downstream machine-learning classifier, we project the same 500 consecutive spikes (**Fig. 4A**) into the four-dimensional feature space (**Fig. 4E**). These 500 spikes represent only a small subset of the full recording, but correspond to the fixed block size later used for model training and testing. Visual inspection of the projected features suggests that the four dimensions are not strongly correlated, indicating that each feature may contribute complementary information about spike morphology or temporal dynamics. Inter-spike interval (ISI): The time duration between the spike and the most recent spike on the same channel (**Fig. 4C**). Clearly, that preceding spike may originate from a different source neuron. The first spike on every channel is excluded. The ISI captures gross firing patterns at the multi-unit level, including bursting behavior.
2. Amplitude: The voltage difference between the trough of the spike waveform and the first ensuing peak (“Peak B”; **Fig. 4D**). We use the term “trough” in this definition, since extracellular electrophysiology studies typically focus on somatic spikes, which exhibit a negative extremum that occurs just before the positive extremum of the intracellular spike. However, we also extract positive spikes, which have a positive extracellular extremum (e.g., **Fig. 3B**). In those cases, the sign of every sample waveform is inverted before deriving the three waveform-based features. Positive spikes have been shown to originate from non-somatic protheses^62^. Therefore, the amplitude feature is signed, corresponding to negative (typically somatic) or positive (typically non-somatic) spikes.
3. Peak A: The (signed) voltage of the nearest peak that precedes the trough (**Fig**. **4D**). A prominent Peak A defines a biphasic spike waveform, which is associated with axonal potentials^62^ (for example, see red unit in **Fig. 4B**).
4. Trough-to-peak time (TPT): The time duration from the extremum (typically, the trough) to Peak B (**Fig. 4D**). The TPT is an easy to compute proxy for the width of the spike, which is often used as one of the features for cell type classification^63,67^.

**Figure 4.**
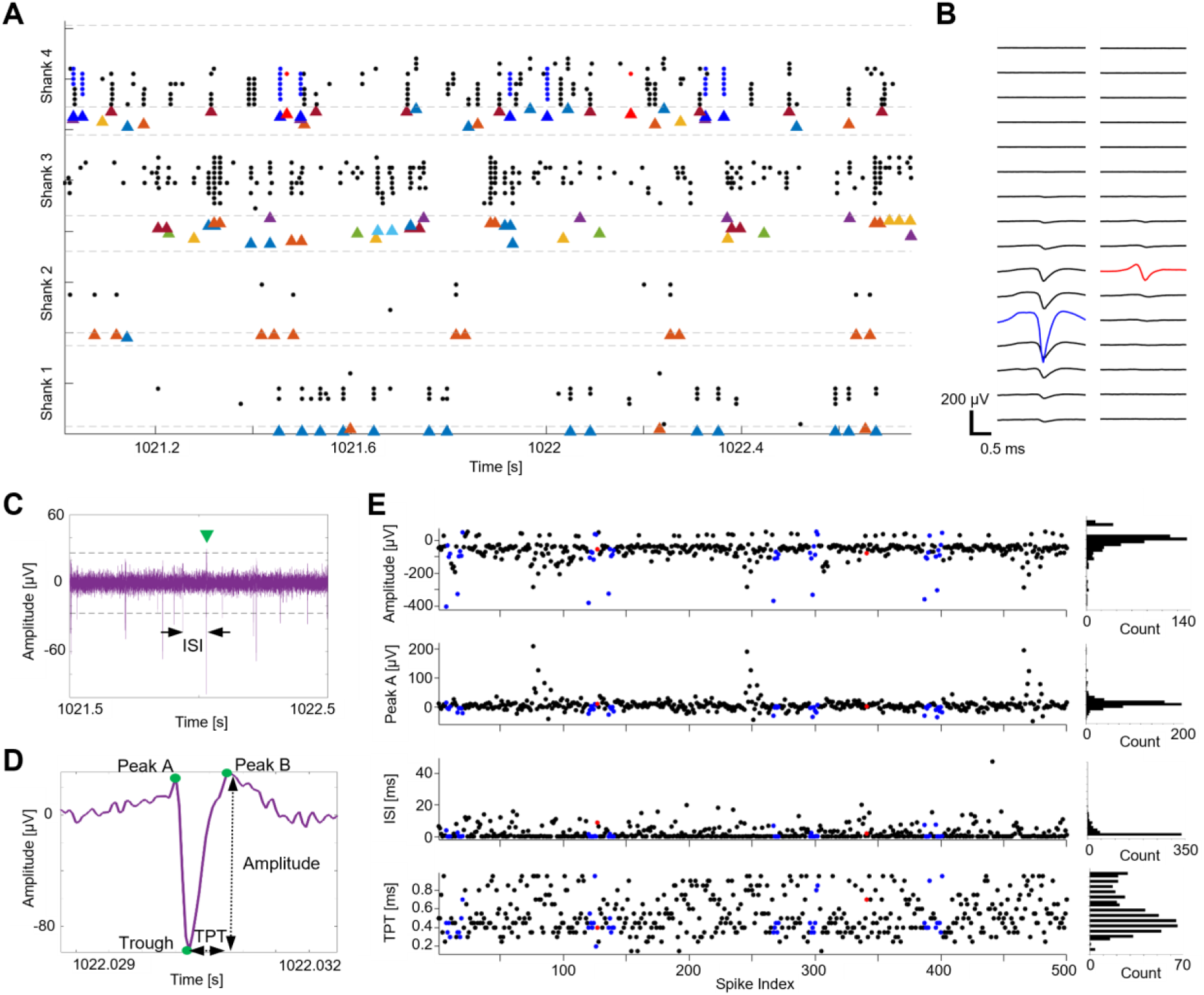
Spike detection and spike feature extraction. (**A**) Raster plot of 500 consecutive, unsorted spikes (circles) and the corresponding single-units (triangles) isolated from the same time window. The raster is divided into four panels corresponding to the four probe shanks. Within each shank, unsorted spikes are plotted according to the electrode depth, where spikes plotted at the bottom row are from the deepest electrode (farthest from the surface of the organoid). Every single-units is shown in a distinct color, time aligned with the unsorted spikes. (**B**) Mean waveforms of two single-units recorded on Shank 4. One single-unit spans multiple channels (blue unit), and the waveforms of the other are confined to a single electrode (red unit). The associated spike detections on Shank 4 are highlighted in (**A**), and their corresponding feature values are shown in (**E**) using the same colors. (**C**) Timing-based spike feature: For every reference spike (green arrowhead), the inter-spike interval (ISI) is defined as the time from the most recent spike (of any unit) on the same electrode. (**D**) Three waveform-based spike features: Amplitude, Peak A, and trough-to-peak time (TPT). (**E**) Feature values of the same 500 spikes and their corresponding distributions shown as histograms (right).

**Figure 5.**
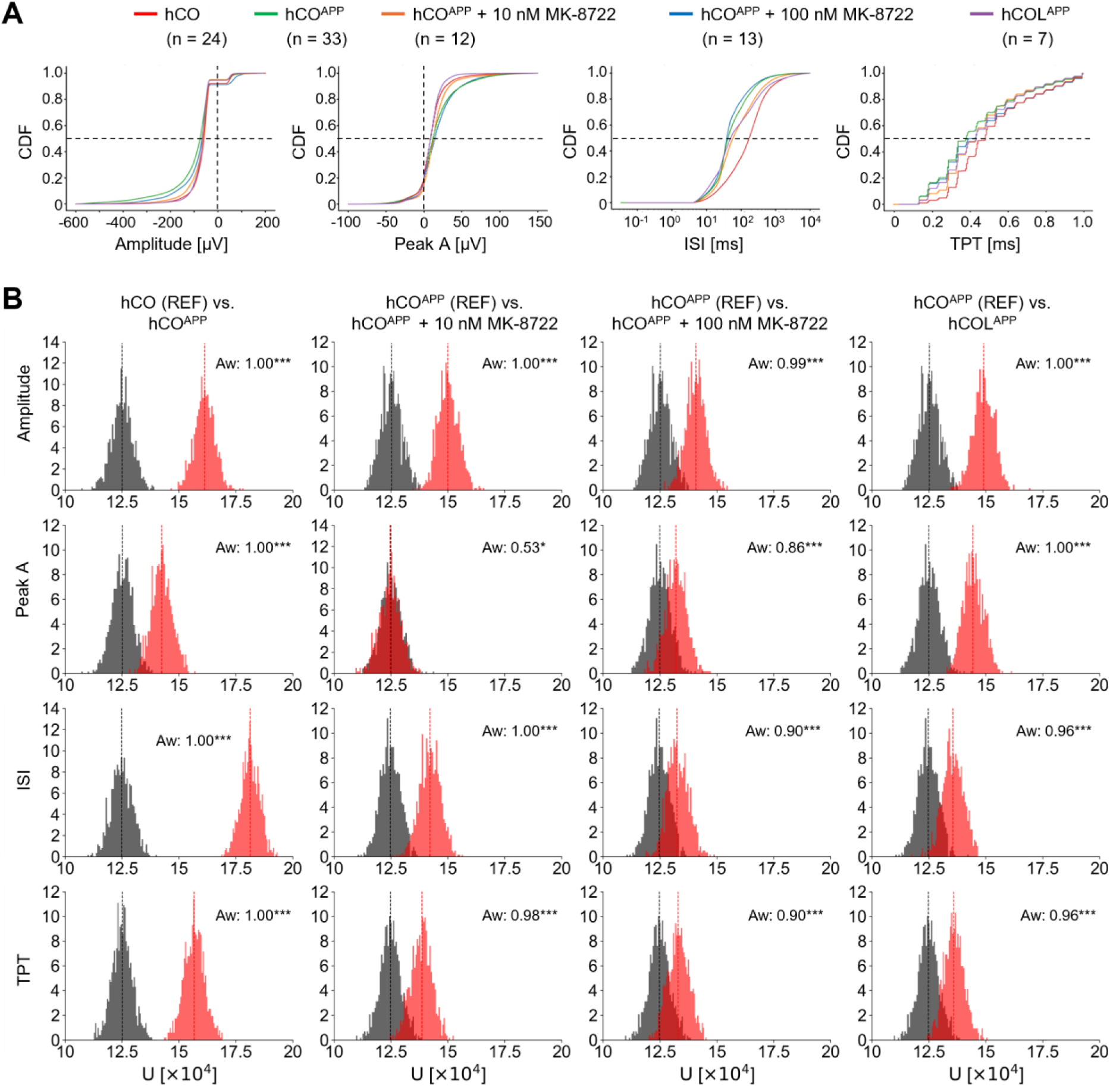
Distributions of spike features and their intra- versus inter-class variability distributions. (**A**) Cumulative distribution functions (CDFs) of spike features for each organoid class, pooled across all recordings. For readability, visualization ranges were clipped for select features. (**B**) Intra- and inter-class feature U-statistic distributions. Each row corresponds to one spike feature, and every column corresponds to a different pair of classes. Gray histograms show the intra-class U-statistic distributions for a reference class (REF), and the red histograms show the inter-class U-statistic distributions, between the reference and the referred classes. Each U-statistic value compares two randomly-sampled groups of 500 spikes from the same class (intra-class distribution) or from two different classes (inter-class distribution). Every histogram is based on 1,000 repetitions, yielding 1,000 U-statistic values. */***: p<0.05/0.001, two-tailed Mann-Whitney U-test.

We extracted and analyzed these four features for each of five organoid classes: hCO, hCO^APP^, hCO^APP^ + 10 nM MK-8722, hCO^APP^ + 100 nM MK-8722, and hCOL^APP^. Overall, we collected data from 89 sessions (89 organoids), yielding 25,815,311 spike events. We computed median and inter-quartile (IQR) statistics for each of the four features of all organoid classes (**Table 1**). Although this set of features does not explicitly characterize spikes as clusters belonging to individual neurons, it captures meaningful differences in the structure and dynamics of neural activity at the individual spike level. For example, the small median ISI value of hCO^APP^ indicates faster spiking compared to hCO, as observed in **Fig. 1**, suggesting that this AD model is hyperactive^68^. The features serve as the foundation for downstream classification in a scalable, automated manner.

### Intra-class feature variability is consistently smaller than inter-class variability

To determine whether the variability of the electrophysiology readout (features) may allow any predictive power of the assay, we first compare spike feature distributions within the same class (intra-class variation) and between different classes (inter-class variation). Intra-class variation reflects differences in structure and circuit dynamics, which may arise from intrinsic (genetic) biological variability, from differences in the environmental conditions during the organoid maturation process, as well as from variability during the data acquisition procedure by an automatically inserted probe. Inter-class variability may arise from the same sources, and from inherent differences between organoid classes. If inter-class variability is consistently larger than intra-class variability, the set of features may provide a useful basis for classification.

To quantify intra-class variability for each organoid class, we pooled and shuffled all spikes from every recording. As neuronal distributions within individual organoids are spatially heterogeneous (**Fig**. **S1**, **Fig**. **S2A**), all class-level statistics were computed by aggregating spike-level features across channels and organoids to minimize the influence of localized sampling variability (**Fig**. **5A**). We then randomly selected two blocks (n=500 spikes each) and computed the Mann-Whitney U-statistic between them. This approach is non-parametric and does not depend on a specific probability distribution. The process was repeated 1,000 times for every feature, generating a distribution of U-statistics (gray histograms in **Fig. 5B**). We chose to compute statistics on small, fixed-size blocks of spikes for two reasons. First, small samples mimic the realistic scenario during high-throughput screening, where only brief stretches of data may be available from each organoid in a large batch. Second, the U-statistic itself depends on the sample size. We used the fixed-size blocks as a building block for training classifiers, ensuring consistency between statistical characterization and predictive modeling. As expected for pairs of similar distributions, the distributions of all intra-class U-statistics were centered around the expected value (n^2^/2=125,000), consistent with no systematic difference in spike features within each organoid class. The spread of the U-statistics across randomly-selected blocks quantifies intra-class variability, serving as a non-parametric reference for evaluating inter-class comparisons.

To assess inter-class variation, we used a similar procedure. For each pair of organoid classes, we randomly sampled 500 spikes from each class and computed the U-statistic between the two blocks. The U-statistic reflects the difference in ranking order between the two samples. This was repeated 1,000 times, generating a distribution of U-statistics for every pair of classes (red histograms in **Fig. 5B**). To quantify the differences, we compared every intra-class (reference) U-statistic distribution to a corresponding (referred) inter-class U-statistic distribution that included the same reference organoid class (gray and red histograms coplotted in the same subpanel in **Fig. 5B**). Quantification was achieved using the Common Language Effect Size (Aw), which estimates the probability that a randomly selected inter-class U-statistic differs from a randomly selected intra-class U-statistic. Aw ranges from 0.5 (completely overlapping distributions) to 1.0 (no overlap), providing an interpretable measure of effect magnitude. The statistical significance (of rejecting the null hypothesis that Aw is 0.5) was tested using a two-tailed Mann-Whitney U-test. Strikingly, all features comparing hCO and hCO^APP^ showed Aw of approximately 1.00 (p<0.001; **Fig. 5B**, first column), suggesting a robust separation between the features of randomly sampled spikes from these two organoid classes. Swapping to hCO^APP^ as the reference class did not alter Aw for all features (**Fig. S5**).

To determine whether the chosen block size of 500 spikes is critical, we conducted a sensitivity analysis using half- and double-sized blocks and repeated the same Mann-Whitney U-test between inter- and intra- U-statistics. Aw increased slightly for all comparisons as the block size increased to 1,000 spike blocks, indicating a greater magnitude of difference in the feature distributions between the class pairs when sample size is larger. Compared with the 500-spike block test, statistical significance obtained from the 250-spike block test was the same for all comparisons (**Fig. S6**); and statistical significance obtained from the 1,000-spike block test was the same for nearly all comparisons, except for Peak A between hCO^APP^ and hCO^APP^ + 10 nM MK-8722 (p<0.001; **Fig. S7**). However, for this specific comparison, Aw increased from 0.53 to only 0.56 when increasing block size from 500 to 1,000 spikes. In summary, at the ranges tested, the feature variability distributions are robust and essentially independent of block size. Statistically significant differences between intra- and inter-class distributions are observed for all four spike features for the organoid class pairs.

### A long-short term memory (LSTM) classifier is trained using blocks of spikes from hCO and hCOAPP

To allow the same probe to sample a large batch of organoids in rapid succession, the assay must produce accurate phenotypic predictions from short electrophysiological recordings, ideally lasting only a few minutes. We developed a machine learning framework that relies on the deterministic, physiologically relevant spike features (**Fig. 5**, **Table 1**) to make binary predictions from brief recordings. Specifically, we trained an LSTM classifier^69^ to distinguish hCO from hCO^APP^ using the spike features extracted from 500-spike blocks.

We split the dataset at the session level. After discarding sessions with less than 5,000 spikes, we assigned 64% of the sessions to training, 16% to validation, and 20% to held-out testing (**Fig. 6A**). Because the number of sessions was limited (**Table 1**), we implemented a block-wise strategy to increase sample size and to increase data variance. Motivated by a previous study^70^, we segmented each session into non-overlapping blocks of 500 consecutive spikes, and every block was treated as an independent sample while preserving the temporal ordering of intra-block spikes. We only retained a maximum of 250 blocks (125,000 spikes) per session to avoid overrepresentation of any session. Therefore, the input to the model was therefore an ordered 500-by-4 spike-by-feature matrix, referred to as a “block”. This strategy increased the number of training samples while constraining the model to learn local temporal patterns, minimizing overfitting and enabling better generalization to novel sessions.

**Figure 6.**
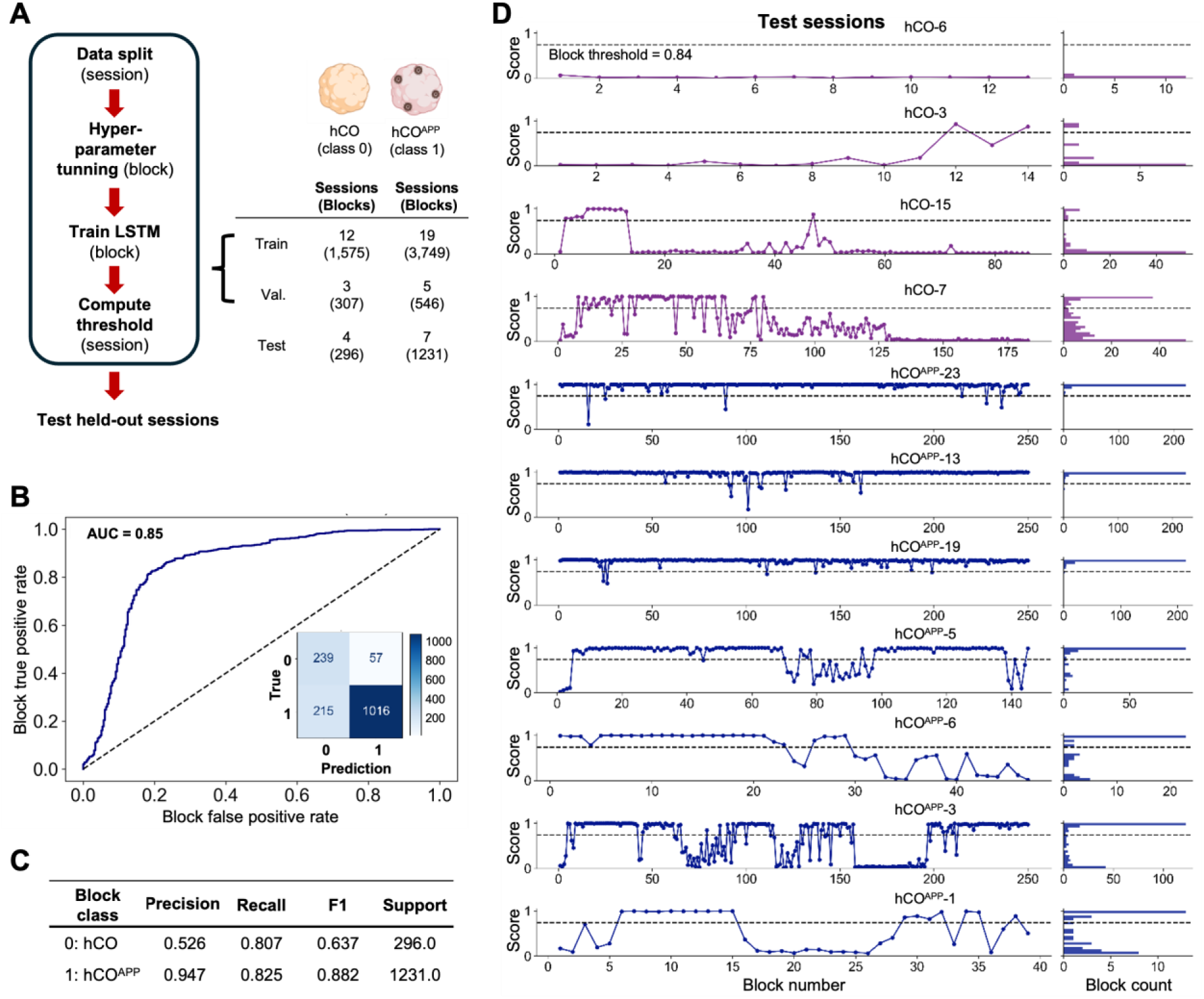
Long short-term memory (LSTM) classifier trained to distinguish hCO and hCO^APP^ spike blocks. (**A**) Overview of model training and testing strategy. The dataset was first split by session, reserving 20% of the sessions for testing. The remaining sessions were split into blocks of 500 consecutive spikes and used to train the LSTM model. A validation split was performed within the training set to guide optimization. During testing, predictions were made at the block level. (**B**) Receiver operating characteristic (ROC) curve for block level predictions on the test sessions. The area under the curve (AUC) reflects the ability of the model to discriminate between classes based on blocks of 500 spikes, independent of a specific threshold. Inset, Confusion matrix for binary classification using the optimal threshold (0.84), highlighting class-specific misclassifications. (**C**) Model classification report for block-level predictions using binary classification showing precision, recall, F1 score, and support for each class. While the model achieved high overall accuracy, precision for class 0 (hCO) was substantially lower (0.53), indicating that many hCO blocks were misclassified as hCO^APP^. (**D**) Block-level prediction scores across test sessions. Left, Scores for each block are plotted chronologically for every hCO (top, purple) and hCO^APP^ (bottom, blue) session. Right, Session-specific histograms of prediction scores illustrate that despite overlap, block distributions are skewed towards their respective class extremes (0 for hCO, 1 for hCO^APP^). Thus, accurate classification may be achieved by aggregating information over blocks.

Block-level predictions were evaluated on spike blocks from the held-out test sessions. At this block level, the model achieved an area under the receiver-operator characteristic (ROC) curve (AUC) of 0.85 (**Fig. 6B**). To assess robustness to dataset partitioning, we performed a post-hoc 10-fold cross-validation at the session level. The set of 57 sessions was randomly partitioned into mutually exclusive folds. In each of ten iterations, a different fold was held out as a test set, while the remaining nice folds were used for training. Model architecture and hyperparameters were the same for all folds. Across folds, the classifiers achieved a median AUC of 0.949 (IQR, [0.720 0.986], indicating consistently strong discriminative performance across different train–test splits.

Using Youden’s J statistic^71^, we determined that the optimal threshold for binary classification is 0.84. Applying this threshold yielded an accuracy of 82%, with precision of 0.95 and recall of 0.83 for the hCO^APP^ (Class 1), and 0.53 precision and 0.81 recall for the hCO (Class 0; confusion matrix in **Fig. 6B, inset** and full classification report **Fig. 6C**). Despite the block level class imbalance (∼4:1, hCO^APP^ to hCO), the model maintains good sensitivity to both classes. To understand how classification errors and successes are distributed within sessions, we plotted the weighted block-level prediction scores for every test session (sorted by class: hCO, **Fig. 6D**, purple; and hCO^APP^, **Fig. 6D**, blue). The block score distributions in every session (histograms shown in **Fig. 6D, right**) revealed that in a few sessions all blocks were classified correctly, appearing at the correct side of the threshold corresponding to the true session label (e.g., **Fig. 6D, top row**). However, other sessions contained blocks which were incorrectly labeled with high confidence. Because the data are inherently heterogenous and because the relevant prediction is at the session (and not the block) level, misclassifications at the block level are both expected and acceptable.

### Session-level classification via confidence-weighted voting distinguishes hCO from hCO^APP^

After training the block-level LSTM model, we derived session-level classification by post hoc aggregation of the block-level model outputs. Specifically, each block classification score 𝑠_𝑖_ ∈ [0,1] was transformed using a confidence-weighting function, 𝑤(𝑠_𝑖_) = 𝑒^𝑐^ ^·^ ^|𝑠^_𝑖_^−0.5|^ where *c* is a scaling constant (set to 4 in this study) that emphasizes high-confidence predictions near 0 or 1. The final session score 𝑦^ was defined as the weighted average of all blocks from that session:

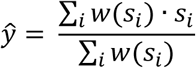

To determine the optimal session-level classification threshold for 𝑦^, we applied Youden’s J statistic to the training and validation sessions, yielding a cutoff of 0.416. We then evaluated model performance on held-out test sessions. Despite the block-level inconsistencies shown in **Fig. 6D**, all test sessions were correctly classified (**Table S2**). Session classification achieved perfect performance on the test set (**Fig. 7A-C**), yielding an AUC of 1.0. Given the small size of the test set, these results should be interpreted as a proof-of-concept for the hierarchical prediction strategy rather than as final performance metrics. The results illustrate how, despite inherent ambiguity at the block level (**Fig. 6C**), aggregating predictions across multiple noisy blocks enables robust session-level classification.

**Figure 7.**
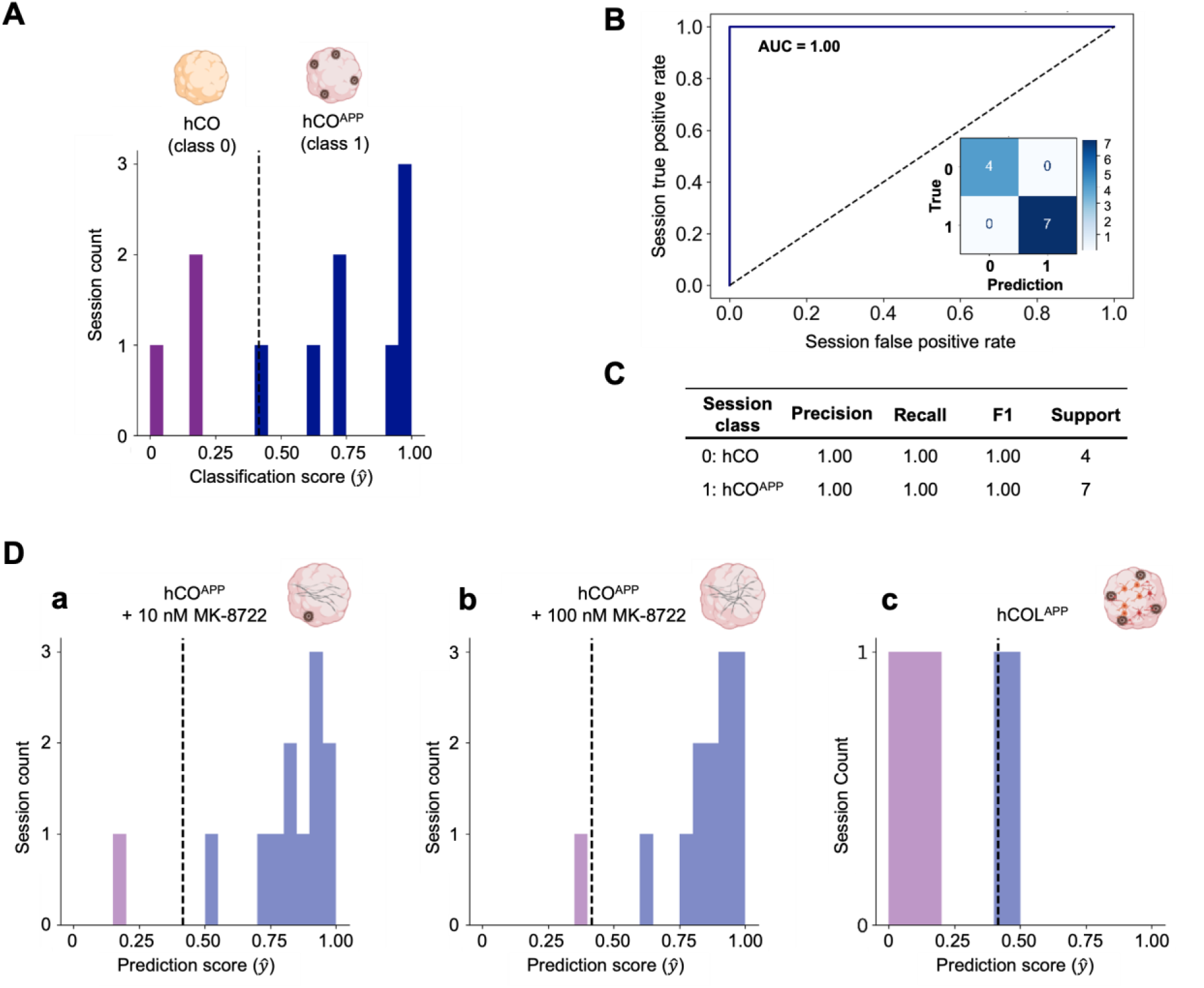
Trained model accurately classifies held-out test sessions and generalizes to novel organoid perturbations. (**A**) Session-level classification scores (ŷ) for labeled test sessions (hCO and hCO^APP^) using a threshold of 0.416 (determined from the training and validation data) yield perfect classification. Histogram bins are colored by their corresponding true class labels (purple for hCO and blue for hCO^APP^). (**B**) ROC for the session-level classification shows perfect performance, with an AUC of 1. (**C**) Classification report details the session-level classification. (**D**) The trained LSTM model is applied to unseen organoid classes to evaluate generalization. **a** hCO^APP^ treated with 10 nM MK-8277 are classified as hCO^APP^ in 11/12 (91.7%) sessions. **b** hCO^APP^ treated with 100 nM of the same drug are classified as hCO^APP^ in 12/13 (92.3%) sessions. Both treatments were applied to the same hCO^APP^ batch and did not result in a detectable functional rescue (classification as hCO), despite the observed reduction in Aβ levels (see Fig. 2D). **c** In contrast, only 2 out of 6 hCOL^APP^ organoids (with enhanced oligodendrocyte differentiation) are classified as hCO^APP^. Histogram bins are colored by their corresponding predicted class labels (light purple for hCO and light blue for hCO^APP^).

### In the hCO^APP^ model, the electrophysiological and molecular phenotypes are inconsistent

To test the utility of the proposed pipeline for capturing the effect of treatments to a disease model, we applied the model to novel organoid classes that were not used during model training. These classes included: hCO^APP^ organoids treated for 30 days with 10 nM or 100 nM of MK-8277, and hCOL^APP^ (**Fig. 2A**). The rationale is that if a treatment induces a detectable change in the electrophysiological phenotype, the classification yielded by the model may shift accordingly. In contrast, if the treatment has no system level electrophysiological effect, classification results would remain consistent with the baseline hCO^APP^ class.

We found that nearly all MK-8277-treated organoids were classified as hCO^APP^. Specifically, MK-8277-treated organoids were classified as hCO^APP^ and not as hCO in 11/12 (91.7%) sessions for the 10 nM group, and in 12/13 (92.3%) sessions for the 100 nM group (**Fig. 7Dab**). These results indicate that under these specific conditions and despite the observed reductions in Aβ levels (**Fig. 2D**), the treatments do not produce a detectable functional rescue.

In contrast, most of the oligodendrocyte-enriched organoids were classified as hCO. Specifically, 4/6 (66%) of the hCOL^APP^ sessions were classified as hCO (**Fig. 7Dc**). Furthermore, even the two sessions classified as hCO^APP^ had prediction scores of 0.4162 and 0.4737. These scores are close to the classification threshold of 0.4161, suggesting a shift towards the control (hCO) phenotype. This result implies that differentiation enriched by oligodendrocyte may have a stronger impact on circuit-level electrophysiology than MK-8277. Clearly, these results pertain to the particular hCO^APP^ model and the specific compound and concentrations used in the experiments, and cannot be used to derive overarching conclusions about BACE1 inhibitors as a drug class.

### High-throughput assay is feasible using data recorded only during probe insertion

To fully exploit the speed and scalability of our electrophysiology platform for high-throughput drug screening, it is essential that the trained models deliver accurate predictions within a minimal recording time. In principle, data collection and model inference can begin as soon as the probe begins to enter the organoid. As described (**Fig. 1**), our system already implements a closed-loop detection algorithm that estimates the high-pass filtered baseline, identifying when the probe electrodes enter the tissue based on the V_RMS_ pre-computed thresholds. A parallel strategy can be implemented for classification: rather than only serving to detect the transition into the organoid, the algorithm can detect spikes, extract features and feed those into a pre-trained classification model to generate block-level predictions as the probe descends in real time. Here, we assess this possibility.

To determine the capability of the system to classify organoids on the fly, we used only the spike data acquired during the insertion phase to evaluate how early and confidently the pre-trained model (**Fig. 6B**) could predict the class of an organoid. These brief datasets, collected during probe insertion (before the probe has reached its final recording depth), were not part of any of the sessions used for model training or testing shown in **Fig. 6A**. The routine begins after a short preparation period involving organoid loading and baseline calibration (**Fig. 8A**). Once the insertion routine is initiated, the probe advances stepwise into the organoid while an algorithm continuously performs spike detection, feature extraction, and block-wise predictions. Early during the insertion, only a few channels are engaged, and block accumulation is initially slow. As more electrodes enter the neuronal part of the organoid, the active channel count increases, spike yield increases, and block acquisition accelerates.

**Figure 8.**
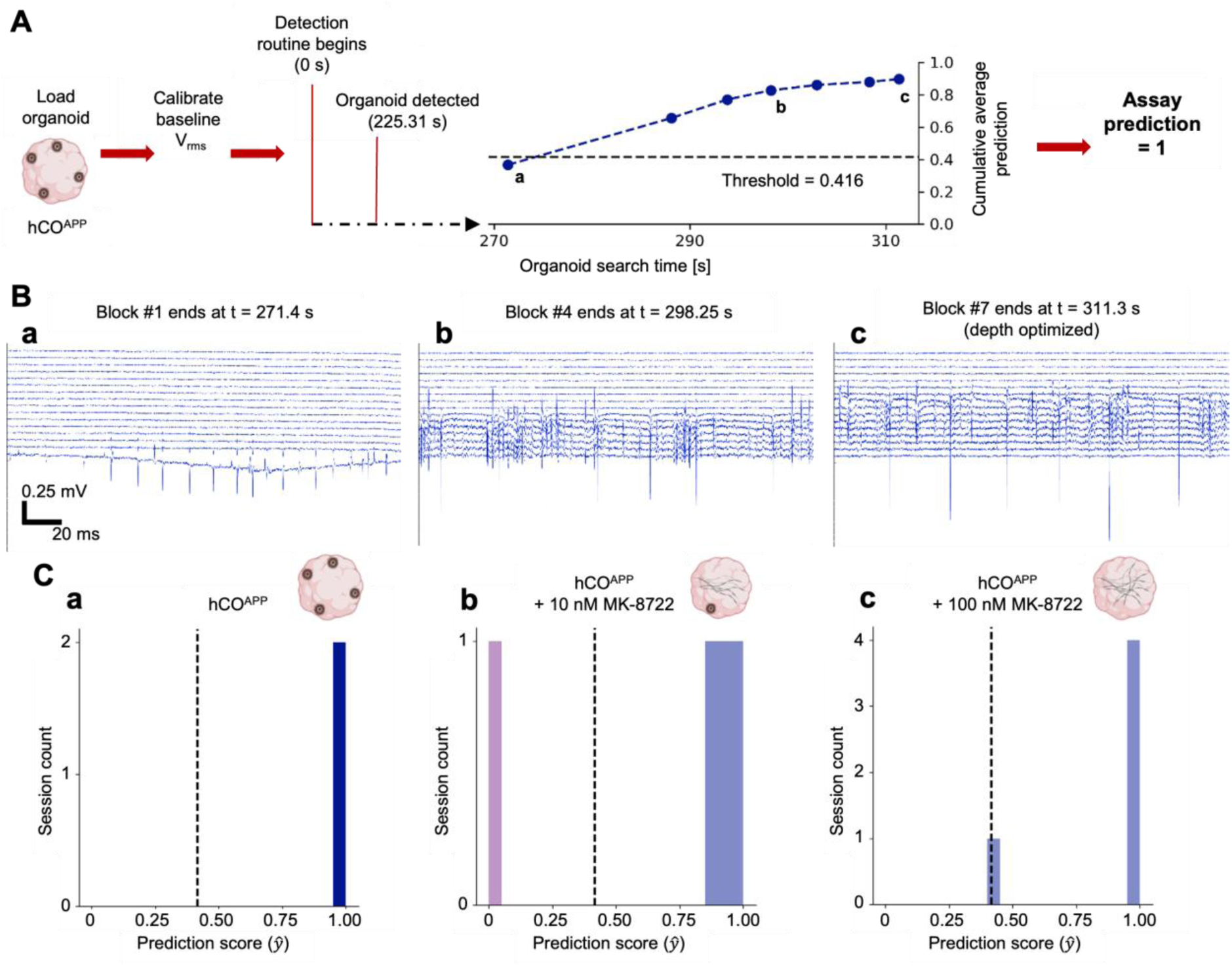
A rapid assay using the pre-trained LSTM model can accurately classify organoid phenotype in real-time during probe insertion. (**A**) Illustration of the automated classification routine. After the organoid is loaded and probe calibration is completed, the detection routine begins (t=0 s). In this example, the organoid is detected at 225.31 s when the first channel exceeds the RMS voltage threshold, and the first spike is detected shortly afterwards. The circles indicate the times at which the acquisition of every 500-spike block has finished. The session level score crosses threshold after the second block (1,000 spikes total) and remains high at the end of the insertion (311.3 s), enabling early and correct organoid classification as hCO^APP^. (**B**) Example wideband recordings (0.2 s windows) captured at the times marked in **A**, illustrating the diversity of neural activity encountered as the probe moves deeper into the organoid. (**C**) Summary of prediction outcomes using insertion-phase data only. **a** 2/2 (100%) hCO^APP^ sessions are correctly classified. **b** 3/4 (75%) of hCO^APP^ organoids treated with 10 nM MK-8722 are classified as hCO^APP^. **c** 5/5 (100%) of hCO^APP^ organoids treated with 100 nM MK87-22 are classified as hCO^APP^.

Representative wideband traces from one probe shank (of four) are presented in **Fig. 8B**, corresponding to the final two seconds of blocks #1, #4, and #7 from **Fig. 8A** (an hCO^APP^ session). These traces illustrate the progressive increase in neural activity as the probe advances into the neuronal region of the organoid. Low-frequency artifacts are occasionally observed during the descent, likely resulting from the mechanical movement of the probe (**Fig. 8Ba**). However, these artifacts do not have a discernible impact on the spikes, which are detected in the high-pass filtered traces (not shown).

To further evaluate whether insertion perturbs transient spiking activity, we quantify spike rates recorded on every channel during the automated insertion (**Fig. S8**). As adjacent electrodes sequentially penetrate the organoid, spike activity typically emerge immediately upon entry into the tissue relative to the media baseline. For electrodes positioned deeper within the organoid, spike rates remain sustained before, during, and after subsequent motor steps, indicating that insertion movements do not significantly alter activity. Occasional brief drops in spike rate are observed only for electrodes located near the organoid surface (Channel 2, at ∼190 s), consistent with minor micromotions that can temporarily displace electrodes out of the organoid at shallow depths. These observations suggest that motor-driven insertion introduces minimal disruption to neuronal activity while enabling progressive sampling across depth. Although electrode motion poses a challenge for stable single-unit isolation, it does not impede feature extraction for our classifier which is based on individual spikes. On the contrary, the dynamic sampling process may enrich the diversity of spike waveforms available to the model as input.

For the hCO^APP^ session illustrated, the first 500 spike block yielded a score below threshold. However, after the second block (1,000 cumulative spikes), the cumulative confidence-weighted average score already crossed the session-level threshold, and then continued to increase as additional data were accumulated (**Fig. 8A**, middle panel). This suggests that reliable classification can be achieved with minimal data under realistic insertion-time conditions.

Finally, we assessed predictions for hCO^APP^ and MK-8277 treated hCO^APP^ organoids that were recorded during the insertion phase (**Fig. 8C**). We found that both (2/2, 100%) hCO^APP^ sessions were correctly classified; that 3/4 (75%) of the 10 nM MK-8277 treated hCO^APP^ sessions were classified as hCO^APP^; and that all (5/5; 100%) of the 100 nM MK-8277 treated hCO^APP^ sessions were classified as hCO^APP^. Overall, probe insertion was completed in a median of 4.43 min (IQR: [4.00 4.63] min), during which a median [IQR] of 7 [1 32] blocks were acquired, each comprised of 500 spikes. The results demonstrate that our automated platform yields rapid and accurate phenotype classification using only spike data collected during the probe insertion phase. This underscores the practicality of the approach for real-time, high-throughput screening.

## DISCUSSION

Accessing functional circuits within 3D brain organoids requires consideration of the spatial decay inherent to extracellular recordings. Planar MEAs have been widely used and are highly effective for surface-level assays. The effective sampling radius of a given electrode is constrained by the physics of extracellular coupling, rather than by electrode technology^72,73^. Specifically, the amplitude of extracellular signals decreases rapidly with distance from the source, limiting sensitivity to neurons located deeper within the tissue. These limitations are increasingly untenable as organoids become larger and more structurally complex^8,16,38,44^.

By inserting high-density silicon probes into intact organoids, the present and prior studies^19,43–45^ overcome the depth limitations and electrode-neuron coupling challenges inherent to planar systems. Originally developed for in vivo recordings, minimally invasive silicon probes provide reliable access to interior layers without disrupting the structure of the organoid^74^. An additional advantage of microfabricated silicon probes is the flexibility of their geometry and site layout, which can be tailored to the anatomical scale and structural heterogeneity of a given preparation. Silicon probes have been extensively customized for diverse in vivo applications, and similar design principles can be applied to organoid systems to optimize channel count, shank length, inter-site spacing, and spatial coverage. The present approach can be extended to support controlled insertion of multiple probes for sampling several distinct regions within the same organoid.

To make deep recordings practical and scalable, we developed an automated system that eliminates user-dependent variability by driving the probe with a motorized stage under real-time feedback control. As the probe advances into the organoid, an RMS-based signal detection algorithm identifies the onset and progression of neuronal activity, halting insertion at the depth that maximizes active channel count. The closed-loop approach ensures consistent and optimized access to the interior, enabling standardized recordings across organoids and users.

This depth-selection strategy is informed by a known biological limitation of whole organoids: diffusion constraints on oxygen and nutrients often produce a stressed or partially necrotic core^47–51^. Experimental measurements have quantified hypoxic length scales of only a few hundred micrometers from the surface, with reduced viability and diminished electrophysiological activity toward the center. This limitation has motivated the development of slice-based preparations^29,75,76^ and more complex vascularization strategies^77^ which can preserve interior tissue viability. Our approach complements these strategies, and is designed to adapt to the prevailing activity landscape by positioning a depth probe where functional neurons are most likely to be active and halting insertion immediately if any probe tip enters a quiet, necrotic region. This strategy is not inherently limited to superficial regions. As organoid engineering advances and healthier, more physiologically sustained interiors become available, the closed-loop approach is expected to enable effective targeting of even deeper neuronal populations. Thus, the system provides a scalable and future-compatible solution for reproducible electrophysiological interrogation of intact organoids across developmental and technological contexts.

In this study, both the MAP2 immunostaining and electrophysiological recording indicate that neurons are enriched within the outer shell of the organoids. Nevertheless, a substantial fraction of the well-isolated single units is detected beyond the immediate surface (**Fig. 3**), which may be missed by surface electrodes. Channels located near the organoid surface on penetrating shanks often fail to detect spikes from deeper neurons, underscoring the need for true depth-resolved access rather than surface or near-surface measurements.

Automated extracellular recording from the organoid interior opens the door to high-throughput functional screening. Importantly, this data acquisition capability must be matched by an analysis framework that supports comparable scalability. Rather than relying on time consuming single-unit analysis, which involves spike sorting and often requires a human expert in the loop, we trained a machine classifier using deterministic features from unsorted spikes.

The features employed in the classification algorithm (e.g., ISI and Amplitude) do not represent properties of well-isolated (single) neurons. Instead, the features are computed from the full detected spike population on each channel, with the rationale of yielding population-level descriptors of network composition and activity. By retaining all detected spikes rather than restricting analysis to well-isolated units, this approach minimizes unit-base selection bias, capturing aggregate dynamics arising from multiple nearby neurons. Because neuronal heterogeneity within an organoid can influence signals recorded by individual electrodes, our classification framework aggregated information across channels and organoids, identifying physiologically meaningful feature differences for robust discrimination while reducing the likelihood that localized variability alone would account for class-level differences.

Our initial results established a reusable and robust framework for binary classification of brain organoids. We employed an LSTM neural network to classify organoid phenotypes based on sequences of spike-level features. The LSTM was chosen over other artificial neural networks (ANNs) due to its ability to model temporal dependencies^69^. For example, two classes may have the same number of isolated versus bursting spikes, but in one class, the bursts may consist of two spikes, whereas in the other the bursts may consist of three spikes. Higher order temporal statistics from individual ISIs may hold circuit-level information essential for classification.

While LSTMs offer advantages, they are also more computationally intensive to train and more susceptible to overfitting in small-sample regimes^69^. In our case, training was performed offline, separately from the experimental workflow, which allowed sufficient time and resources for model development. Once trained, the classifier can be used in many new experiments, supporting real-time classification without retraining. As data acquisition is scaled up, the risk of overfitting is expected to decrease. We acknowledge that a simpler ANN might have been sufficient for the current binary task, but this was not tested exhaustively. Looking forward, more challenging classification problems such as subtle drug responses or complex genetic models may also benefit from the temporally aware architecture of the LSTM classifier, which is aligned with the temporally dynamic nature of brain function and circuit-level computations^78^.

Feature-wise statistical comparisons (**Fig. 5B**) and binary classification outcomes (**Fig. 7**) probe different properties of the data and are not expected to align perfectly with one another. The U-statistic analyses are univariate and sensitive to subtle distribution shifts in pooled spikes. In contrast, the binary classifier integrates information over multiple features while accounting for temporal structure and is evaluated at the block and session levels. Consequently, a treatment may induce detectable shifts in individual features without producing a separable phenotype under the multivariate patterns used for classification.

As a proof of concept, we generated cortical organoids from hiPSCs with and without Swedish/Indiana mutations in the APP gene, mutations widely used in familial AD modeling due to their effect on Aβ overproduction. The resulting hCO and hCO^APP^ lines formed the basis for the binary classification task, serving as an initial benchmark for validating the functional screening framework. For simplicity, binary classifiers are often sufficient for functional assays, where the objective is to determine whether a compound elicits a specific phenotype, such as therapeutic efficacy or toxicity^79^. The same feature extraction and LSTM-based framework is extensible to multiclass settings, enabling more refined classification when warranted. Indeed, our primary goal in this study was not to optimize the classification of a specific disease model, but rather to establish a framework that generalizes to other applications, including assessing genetic mutations, pharmacological perturbations, or environmental toxicants^80^. For example, the U.S. Environmental Protection Agency’s ToxCast program has identified thousands of uncharacterized neurotoxicants that remain unscreened^81^.

Our model was trained on short blocks of 500 spikes, which were derived from longer continuous sessions. This design allowed for consistent input dimensions and high training sample counts^70^. However, it introduced a challenge: spike blocks were only weakly labeled (inherited from the parent session) and may not uniformly reflect the session phenotype. Variability in organoid maturation, spontaneous circuit states, and random sampling by the probe introduced considerable heterogeneity across blocks. As expected, some blocks yielded confident scores aligned with the session label; others hovered near chance or even misaligned with the session class. This is not a model failure, but rather a reflection of real-world variance. Instead of discarding these fluctuations, we exploited them using a confidence-weighted aggregation strategy, amplifying the influence of high-confidence blocks. This can be especially useful to separate two classes with identical mean feature values, but one would be more variable over blocks than the other.

In one experiment, treatment with a BACE1 inhibitor (MK-8722) reduced Aβ levels in hCO^APP^, as confirmed by immunostaining, yet the electrophysiology-based model continued to classify the treated organoids as AD-like. Because the hCO^APP^ line carries the APP Swedish mutation, which has been reported to show reduced sensitivity to BACE1 inhibition, we did not expect complete normalization of Aβ production or electrophysiological phenotypes. Instead, this experiment was designed as a pharmacological probe to test whether our platform can detect a directionally appropriate modulation of Aβ-related pathology^82–84^. The observed effects pertain to the specific mutation, background, compound, and concentration. However, the discrepancy between changes in Aβ immunostaining and persistent AD-like network activity underscores the value of a functional readout. This distinction is particularly relevant for neurodegenerative diseases (e.g., AD), where molecular hallmarks may not map directly onto behavioral or physiological outcomes^57,58,85^. Complementary to quantifying the biochemical state, our assay measures functional activity at the circuit level, which underlies all brain functions and dysfunctions.

Although the mechanism underlying AD pathology is not explored in this study, the elevated spiking activity in APP organoids is consistent with prior reports of network hyperactivity and excitatory-inhibitory imbalance in AD-related models^68^. In the future, post-hoc interpretability methods^86^ may be applied to quantify the relative contribution of individual features to prediction performance.

To push the limits of throughput, we evaluated whether classification could be achieved using only data collected during the probe insertion phase, without requiring extended stationary recording. This test revealed that the model could accurately classify sessions using just a few minutes of insertion-phase activity. As the probe descended and encountered more active neurons, block accumulation accelerated and predictions converged rapidly. This insertion-only mode points toward true real-time screening applications, where the classifier operates concurrently with data acquisition, which is in turn synchronized with the instrument robotics. Block-level misclassifications can occur in sequences of multiple blocks, likely due to the intrinsic temporal correlations of neuronal activity and possibly due to transient changes in network states. Aggregating predictions across multiple blocks remains effective because the erroneously-classified stretches are brief compared with the correctly-classified blocks.

While this study demonstrates the feasibility of automated, single-organoid targeting, the platform presented is readily extensible to multi-organoid probing via X-Y actuation, allowing the probe to access multiple wells within the same apparatus. Although a single probe may eventually be designed to target multiple wells simultaneously, full-plate coverage is unlikely due to limited depth optimization, noise accumulation, system complexity, and cost. Thus, it is essential that useful information will be acquired rapidly, minimizing probe dwell time per well. The feasibility of this scalability is already evident from the results reported in the present study: because the full workflow completes within minutes, sequential probing across wells is possible.

Looking ahead, a key feature of this platform lies in its ability to support the cumulative development of a growing portfolio of pre-trained classifiers. As we expand the range of tested compounds, disease models, and experimental conditions, the resulting database can be used to train increasingly accurate and generalizable models tailored to specific toxicants, genetic mutations, or therapeutic targets. ^87^We expect that over time, this resource will enable plug-and-play classification across new assays. Continuous addition of new sessions supports iterative refinement, transfer learning, and cohort-level aggregation, paving the way for robust, reproducible, and scalable phenotyping across diverse organoid systems.

## ACKNOLWEDGEMENTS

This work was partially funded by Merck Sharp & Dohme LLC, a subsidiary of Merck & Co., Inc. We thank Danny Bitton, Michael Wurst, and Jakub Arnold from MSD Prague for helpful discussion on machine learning model.

## AUTHOR CONTRIBUTIONS

Conceptualization: R.M., B.C., B.J., E.S.; Instrument design: F.W., L.A., B.J.; scRNAseq library generation and analysis: S.K.V., V.M.P., A.M.T, B.C.; conceptualization of AD disease modeling: M.E.K., I.K., B.C; Data acquisition: F.W., L.A., B.C.; Electrophysiology analysis: F.W., E.S.; Machine learning framework: F.W., C.S.; Writing: F.W., E.S., B.J., B.C.

## DATA AVAILABILITY

The data that support the findings of this study are available from the corresponding author upon reasonable request.

## METHODS

This study complies with all relevant ethical regulations approved by Merck Research Laboratories (MRL). All experiments involving human iPSCs were approved by the Merck Stem Cell for Drug Discovery (SCDD) Committee. The human brain tissues assessed in this study were obtained from Dx Biosamples LLC (Sample ID # 531888A(3), San Diego, CA) based on MRL Human Investigation Committee. It has been approved for use with a waiver of consent.

### Human iPSCs culture

XCL-1, XCL-1_APP Ind (SwHomo)-3G05-E01, and BC1 hiPSCs were cultured on Matrigel (Cat # 356230, Corning, NY) coated cell culture dishes with mTeSR1 plus media (Cat #100-0276, STEMCELL Technologies, BC, Canada). hiPSCs were passaged every week by treatment with Dispase (0.83 U/ml, Cat # 07913, STEMCELL Technologies, BC, Canada).

### Generation of hCO variants

As described earlier^53^, we generated hCOs and hCO^APP^ by using XCL-1 and XCL-1_APP Ind (SwHomo)-3G05-E01 human iPSCs, respectively. Briefly, after dissociating cells via Accutase, a total of 9000 cells were plated into a well of U-bottom ultra-low-attachment 96-well plate in neural induction medium (DMEM-F12, 15% (v/v) KSR, 5% (v/v) heat-inactivated FBS (Life Technologies),1% (v/v) Glutamax, 1% (v/v) MEM-NEAA, 100 µM β-Mercaptoethanol) supplemented with 10 µM SB-431542, 100 nM LDN-193189, 2 µM XAV-939 and 50 µM Y27632). FBS and Y27632 were removed from day 2 and 4, respectively. The medium was replenished every other day until day 10, where organoids were transferred to the ultra-low-attachment six-well plate. The organoids were cultured in spinning hCO, dorsal patterning (day 10 to day 18) medium with minus vitamin A (1:1 mixture of DMEM-F12 and Neurobasal media, 0.5% (v/v) N2 supplement, 1% (v/v) B27 supplement without vitamin A, 0.5% (v/v) MEM-NEAA, 1% (v/v) Glutamax, 50 µM β-Mercaptoethanol, 1% (v/v) Penicillin/Streptomycin and 0.025% Insulin). The dorsal patterning medium was replenished every other day until day 18, when media was switched to the maturation media, hCO medium with vitamin A (the same composition as described above except B27 with vitamin A) supplemented with 20 ng/ml BDNF and 200 µM ascorbic acid. The maturation medium was changed every 3 days after day 18.

To generate AD forebrain variants, dorsal and ventral forebrain organoids were developed using XCL-1_APP Ind (SwHomo)-3G05-E01 and 3-TF-expressing BC1-iPSCs. AD control organoid variants utilized 100% XCL-1_APP Ind (SwHomo)-3G05-E01) while AD variants with OL-lineage cells were created by mixing 20% BC1-iPSCs, driving OL-lineage cells, and 80% non-infected XCL-1_APP Ind (SwHomo)-3G05-E01) and followed by dorsal patterning and maturation.

### Automated multiplex immunofluorescence staining

Organoids were collected in a 6-well plate and fixed in 4% paraformaldehyde (PFA) for 24 hours at 4 °C as described previously^53^. Following fixation, single organoids were transferred to individual cassettes for paraffin embedding. The tip of a tube was placed on the embedding mold, and paraffin wax was melted; once melted, organoids were carefully moved into the mold^87^. Each paraffin block contained three organoids per condition, along with AD patient brain tissue. The blocks were sectioned at 5 µm thickness onto charged glass slides, allowing for serial sections to be obtained from nine organoids across three different conditions. For immunofluorescence staining, slides were baked at 60 °C for 1 hour, deparaffinized using the BOND Dewax protocol, and subjected to heat-induced epitope retrieval with BOND Epitope Retrieval Solution 2 for 20 minutes. Staining was performed on the Leica Bond RX stainer using BOND Ready-To-Use primary antibodies of monoclonal mouse or rabbit origin. The immunofluorescence protocol (Opal) was executed sequentially, with chromogens mixed on-board the BOND RX as part of the automated staining process, utilizing enzymatic conversion with BOND Polymer Refine Detection (HRP) and BOND Polymer Refine Red Detection (AP). Finally, images were scanned at 20x magnification using the Vectra Polaris and unmixed for analysis. Slides were incubated in primary antibodies (anti-AT8 1:100, anti-MAP2 1:100, anti-6E10 1:100, anti-SOX2 1:200, anti-Ki67 1:1000, anti-vGAT 1:100, anti-vGLUT2 1:100, anti-OLIG2 1:200, anti-MBP 1:1000).

### Aβ-ELISA

Aβ peptide levels were quantified using commercially available sandwich ELISA kits (human Aβ40, Invitrogen KHB3481; human Aβ42, Invitrogen KHB3441). Organoids were homogenized, and total protein concentration in 20% (w/v) homogenates was determined using a BCA assay. An aliquot containing 200 µg of protein in 400 µl was ultracentrifuged at 100,000 × g for 1 h at 4 °C (L100K ultracentrifuge, Beckman-Coulter, Brea, CA). The resulting supernatant was collected as the soluble fraction. The pellet was resuspended in 50 µl of 70% formic acid, sonicated, and then neutralized with 2.5 M Tris buffer, pH 8.5, to obtain the insoluble fraction. Aβ40 and Aβ42 levels were measured in both soluble and insoluble fractions according to the manufacturers’ protocols. Absorbance was recorded at 450 nm using a microplate reader (EL800, BioTek, Winooski, VT), and concentrations were calculated from standard curves supplied with the kits.

### Library preparation of scRNAseq

Cortical organoids 50 and 115 day old were randomly collected from 3 different culture dishes, with 6-10 organoids pooled together. As previously described^38^, the organoids were initially dissociated using the papain dissociation system according to the instructions of the manufacturer (Worthington Biochemical Corporation, NJ). Subsequently, after washing with Hank’s balanced salt solution, the organoids were dissected into small pieces in papain solution and oxygenated with 95% O_2_:5% CO_2_ for 5 minutes, and then incubated at 37 °C for 1 hour. Following the generation of a single-cell suspension via trituration, the single cells were suspended in 1% BSA/PBS supplemented with 10 µM Y27632 and stained for Aqua Dead Cell stain (Cat. No. 50-112-1525). FACS sorted live cells were re-suspended in 0.04% BSA/PBS (128 cell/µl) and used to generate cDNA libraries by utilizing the Single Cell 5’ Reagent Kits on a Chromium-X platform. In brief, the cells were partitioned into nanoliter-scale Gel Bead-In-Emulsions (GEMs), and microfluidic cells were streamed at limiting dilution into a stream of Single Cell 5’ Gel Beads and then a stream of oil. After cell lysis, primers, an Illumina P7 and R2 sequence, a 14 bp 10xBarcode, a 10 bp randomer, and a poly-dT primer were released and mixed with the cell lysate and a bead-derived Master mix. These cDNAs were used to generate sequencing libraries for Illumina platforms using library generation kit according to manufacturer supplied user guides (10X Genomics, CA). The libraries were sequenced using the Illumina Novaseq 6000 platform.

### Data processing of scRNA-seq

scRNA-seq FASTQ files were processed and generated count matrix over human reference version hg38 using CellRanger (v 8.0.1) with default parameters. Before processing scRNA-seq analysis, we confirmed the low doublet frequency of our scRNA-seq libraries (mean±SEM, 0.82±0.28%) by counting cells expressing both TBR1 and GFAP, which are usually exclusively expressed in cortical neurons and astrocytes, respectively.

The scRNA-seq libraries from forebrain organoid variants and human fetal brains^88^ were collectively analyzed using Seurat (v5.0) within the R (v4.4.1) environment. Initially, as a part of quality control measures, cells with fewer than 1,000 detected genes and genes expressed in less than 5 cells were excluded. For each organoid study, the feature counts were normalized to the total counts and multiplied by a factor of 10,000. Highly variable features (genes) were then selected using variance stabilizing transformation. Following the prioritization of features based on their variance across scRNA-seq libraries, the top 2,000 features were retained for downstream analysis. Cell pairwise anchor correspondences between different single-cell transcriptome studies were identified using 30-dimensional spaces from canonical correlation analysis. These anchors were then utilized to integrate scRNA-seq datasets and transform them into a shared space. Gene expression values were scaled for each gene across all integrated cells and used for principal component analysis. Subsequently, using the first 30 principal components (PCs), all cells were projected onto a two-dimensional UMAP space. Graph-based cell clustering was implemented with the shared nearest neighbor method from the first 30 PCs and a 0.5 resolution value. Differentially expressed genes in each cluster were identified by employing a threshold of more than 1.25 average fold change and p<0.05 through a two-sided t test. Overrepresented gene ontology terms were identified using GOstats (v2.24.0), and the false discovery rate (FDR) was estimated using the Benjamini-Hochberg method with the p.adjust function. An FDR below 0.05 was used as a threshold for statistical significance.

Cell types were assigned to the clusters with slight modifications from a previous method^53^. Initially, the clusters were categorized into two main groups, neuronal and non-neuronal, based on general neuronal markers (STMN2, GAP43, and DCX) and early neurogenesis genes (VIM, HES1, and SOX2). Within the neuronal group, further classification into cortical neurons (CN) and interneurons (IN) was conducted based on the expression of specific markers, such as TBR1 and amino acid synthetic enzymes and transporters (GAD1, GAD2, vGLUT1, and vGAT). Additionally, glutamatergic and GABAergic neurons were identified by the expression of their transporter genes (SLC17A7, SLC17A6, and SLC32A1), while neuronal clusters lacking these specific genes were annotated as immature neurons.

Cell clusters expressing biglycan and decorin were allocated as proteoglycan-expressing cells (PGC). Furthermore, clusters enriched with genes related to “Cilium assembly (GO:0044458)” and “myofibril genes” (GO:00300016) were labeled as cilium-bearing cells (CBC). Clusters exhibiting significant overrepresentation of “Glia cell differentiation (GO:0010001)” were classified as glial cells and further annotated as oligodendrocyte progenitors (OPC), oligodendrocytes (OL), and astrocytes (AS) based on unique markers for each cell type (PDGRFA, PTPRZ1, BCAN, OLIG1/2, PLP1, GFAP, SLC1A3, and S100B) and relevant GO terms. The remaining cluster was designated as intermediate.

To validate the cell type annotation, gene signatures of neuron, astrocyte, and OPC obtained from single-cell transcriptome in human fetal and adult brains^89^ were used to pre-rank genes by relative expression and evaluated using GSEAPY software, as described previously^54^.

To compare the cell types in our study across different time points with those in established organoid protocols from the literature, we utilized transcriptomic data from the Human Neural Organoid Cell Atlas (HNOCA)^55^. We accessed the publicly available section of HNOCA, which encompasses all integrated protocols except for three that remain unpublished. We employed the CellHint tool^90^ v0.1.1 to harmonize cell types across various organoid protocols and to compute a cross-dataset distance matrix. This matrix included 1,665,880 cells and 161 cell types from 26 different protocols, as indicated in the “assay differentiation” section of the HNOCA metadata. During the harmonization process with CellHint, each cell was assigned to the most similar cell type from each dataset, resulting in a comprehensive assignment matrix. The distance score in this matrix quantifies the transcriptomic dissimilarity between cell types across datasets, computed as the cosine distance between their gene expression profiles, with higher values indicating greater differences^55^. For ease of analysis, we focused on specific comparisons: excluding all cell types labeled as ‘unknown’ and retaining only those comparisons that involved our datasets and had a distance score greater than 0.5 for at least one cell type. Finally, we visualized the inferred similarities among cell populations using a UMAP plot, which displayed unsupervised hierarchical clustering

### Instrument design and operation

We designed an automated system to precisely drive a microfabricated silicon probe into stationary brain organoids. The core sensor is a 64-channel silicon probe (P64-1, Diagnostic Biochips Inc., Glen Burnie, MD) connected to a custom printed circuit board housing a front-end ASIC (RHD2164, Intan Technologies, Los Angeles, CA). This assembly interfaces with a data acquisition system (Open Ephys, Lisbon, Portugal) via SPI communication. The probe and ASIC are mounted on a Z-axis micromanipulator (KMTS50E, Thorlabs, Newton, NJ), which is computer-controlled. A custom Python script asynchronously controls the Open Ephys system to start/stop recordings and reads recent data to compute RMS voltage for organoid detection. Upon detection, the system halts further probe advancement.

### Data acquisition

Recordings were performed with the system set inside an incubator maintained at 37 °C and 5% CO₂ (95% air) (Hercell vios 160i, ThermoFisher, Waltham, MA). Organoids were transferred manually from their culture plate into a recording well using a pipette. The well was preloaded with 0.3 ml of warm BrainPhys medium and topped off with another 0.5 ml post-transfer. After loading the well and the probe into their respective mechanical fixtures, the center of the probe tips (midpoint between shank 2 and shank 3) was automatically aligned to the center of the well by design. A custom Python script then controlled the Z-axis actuator to insert the probe 2 mm above the bottom of the well to provide baseline RMS measurements. The baseline measurement depth was configured based on predetermined maximum size of the organoids. Automated descent proceeded at steps of 50 µm, until one or more channels became active, at which point step size was reduced to 20 µm for a more precise depth control. The insertion stopped if any of the following conditions were met: (1) the probe reached the bottom of the well; (2) no new active channels were detected during three consecutive steps; or (3) the leading channel near the tip of any of the shanks became inactive (fell below the pre-computed threshold). This closed-loop, precision driving helped to avoid overshooting into necrotic or immature zones. Recordings were acquired for at least one hour at a sampling rate of 20 kHz, corresponding to approximately 9 GB/hour.

### Single-unit analysis

Spike sorting was performed on all recording sessions using Kilosort4^66^. We isolated 1,358 single-units that were labeled by Kilosort4 as “Good Units” and had ISI index^91^ below 0.2.^63^.

### Spike feature extraction

A custom Python algorithm processed raw recordings by applying a bandpass filter (300-6,000 Hz), followed by spike detection using a 5 SD threshold. Each detected spike waveform consisted of 40 sample points spanning a 2 ms window centered at the trough (maximum absolute voltage deflection). We computed four spike features (**Fig. 4**) for every spike and extracted spike-level metadata, including timestamp, channel, and session ID. These were stored in concatenated Pandas DataFrames^92^, grouped by organoid class and encompassing all spikes from corresponding sessions.

### Binary classification of spike blocks using an LSTM Model

We split all hCO and hCO^APP^ recording sessions into training (64%), validation (16%) and testing (20%) sets. The LSTM model was trained and optimized using the training/validation sets and all performance metrics were measured using the testing set. Since a limited number of sessions was available, we segmented each session into 500-spike blocks as inspired by a previous work^70^, in order to increase data variance. Each block was treated as a separate training instance while maintaining intra-block temporal order. To avoid overrepresentation of long sessions, we retained a maximum of 250 blocks (first 125,000 sequential spikes) per session. The resulting model input was a 500 by 4 spike-by-feature matrix for every block. Separate files tracked metadata associated with every block, including unique session identifier, class label, and split category (train/validation/test).

We implemented the LSTM classifier using TensorFlow/Keras. We used Optuna^93^ to optimize hyperparameters including the number of recurrent units, dropout rate, dense layer width, and learning rate. The best-performing architecture comprised an LSTM layer with 64 input units, dropout rate of 0.1, a 32-unit ReLU dense layer, and a single sigmoid output unit. Training was performed using binary cross-entropy loss and the Adam optimizer^94^ (learning rate = 0.001).

### Statistical analyses

All statistical tests used a significance threshold of α=0.05. Descriptive statistics (n, mean, median, IQR, SD, SEM) are reported in the text, figure legends, and tables. Group medians were compared using two-tailed Mann-Whitney U-tests. Statistical significance is annotated in figures as follows: ns: p>0.05; */**/***: p<0.05/0.01/0.001.

