## Supplementary Figures for "High-density extracellular recordings from the interior of intact brain organoids enable automated high-throughput functional assay"

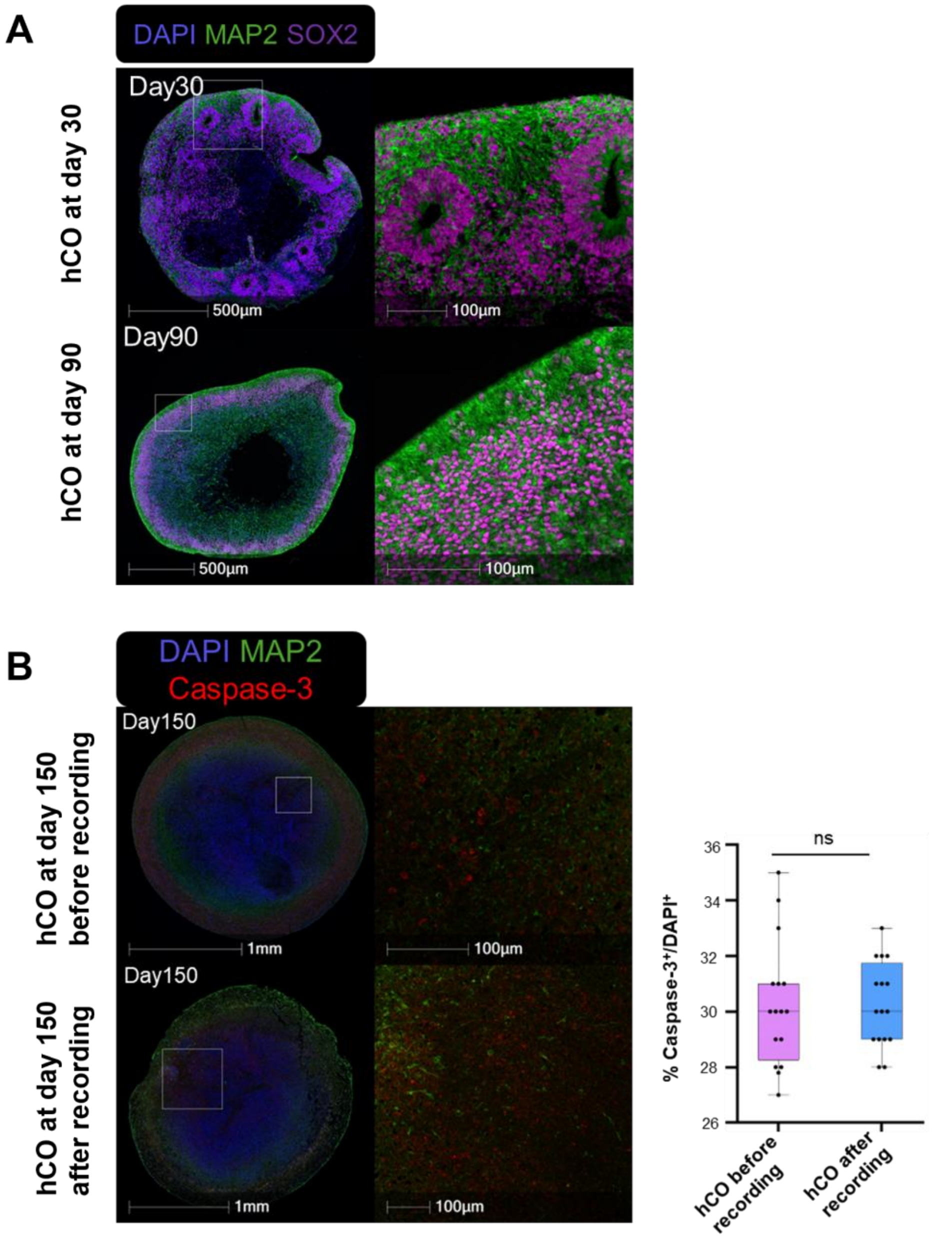


**Figure S1. Radial maturation of neuronal layers in human cortical organoids (hCO) and preserved neuronal health after acute recording**. (**A**) Immunofluorescence staining of hCO at day 30 and day 90 using MAP2 (neuronal marker) and SOX2 (neural progenitor marker). (**B**) Acute recordings do not increase cell death in day-150 organoids. Representative immunostainings of day-150 organoids fixed before (top) or after (bottom) a 1-hour recording session in the device, showing DAPI (blue), MAP2 (green), and cleaved caspase-3 (red). Radial maturation pattern of the neuronal shell is preserved after acute recording. Higher-magnification views of the boxed regions are shown in the middle panels. Right, quantification of cleaved caspase-3–positive cells per unit area does not show significant difference between pre- and post-recording organoids (n = 16 organoids per group; ns=0.74226, Mann–Whitney U-test), indicating that probe insertion and 1-hour recordings do not measurably increase apoptosis under our conditions. Scale bars: 1 mm (left), 100 µm (middle).

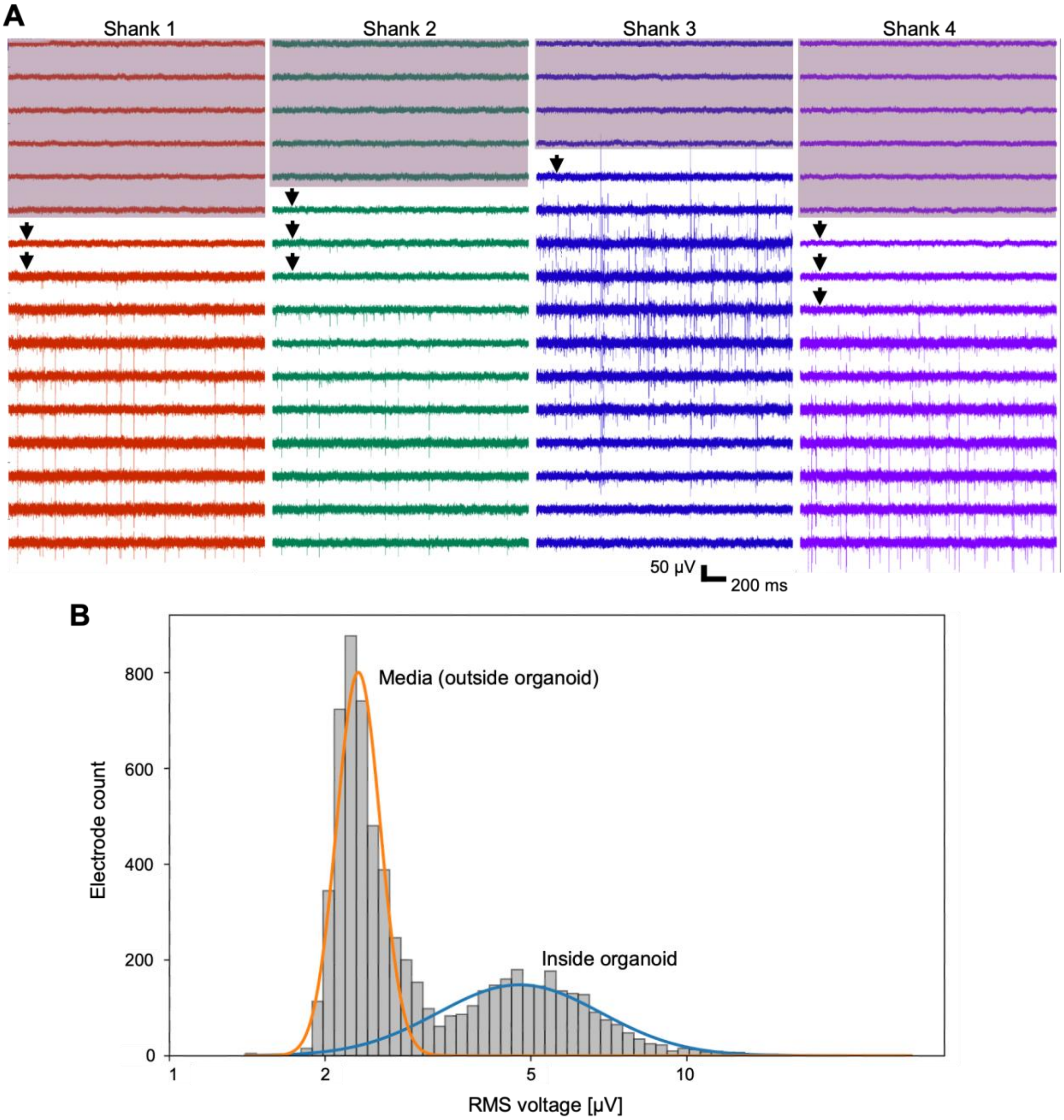


**Figure S2. Electrodes inside or outside of an organoid can be distinguished by $V_{RMS}$.** (**A**) Example two-second, 64-channel recording from an hCO. Each column represents band-pass filtered (300-6,000 Hz) traces recorded by one probe shank with 16 recording channels, sorted according to their physical depths: the bottom-most trace is deepest. Electrodes are separated by 20 µm in depth. Shaded channels (top) are outside the organoid. Non-shaded channels (bottom) are inside the organoid as determined by the RMS voltage threshold computed for each channel. Channels inside the organoid but close to the surface (arrowheads) may not detect spikes (5 SD above the mean $V_{RMS}$). (**B**) The RMS voltage values from 6,656 electrodes across 104 recordings are fitted with a two-component one-dimensional Gaussian Mixture Model. The component corresponding to electrodes in the media (outside the organoid) has a mean log10(RMS) of 0.366 and SD of 0.042 (n=4,073; weight=0.591; orange). The component corresponding to electrodes inside the organoid has a mean of 0.677 and an SD of 0.157 (n=2,583; weight=0.409; blue).

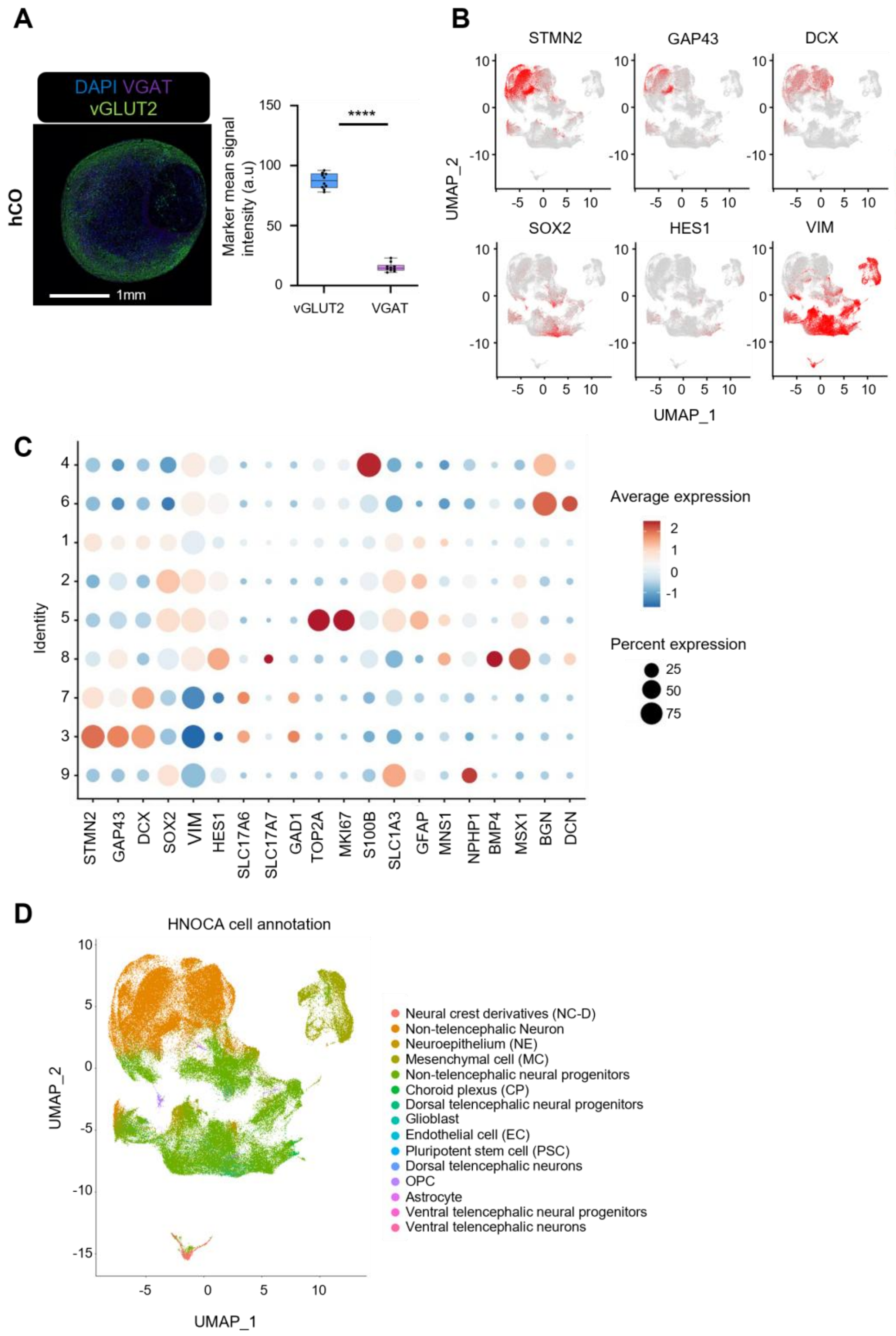


**Figure S3. Characterization of hCOs**. (**A**) Left, Immunostaining of an hCO at day 120 for the glutamatergic neuron marker vGLUT2 and the GABAergic neuron marker VGAT. Right, Quantification of markers. Box plots show medians and IQRs over three independent batches (n=10 organoids; ****: p<0.0001, two-tailed Mann-Whitney U-test). (**B**) Expression patterns of genes related to early neurogenesis and neuronal growth cone. (**C**) Comparison of gene markers related to lineage commitment among clusters. (**D**) Cells from our datasets were projected to the Human Neural Organoid Cell Atlas (HNOCA)[38] using its computational tools, confirming the presence of cells relevant to forebrain.

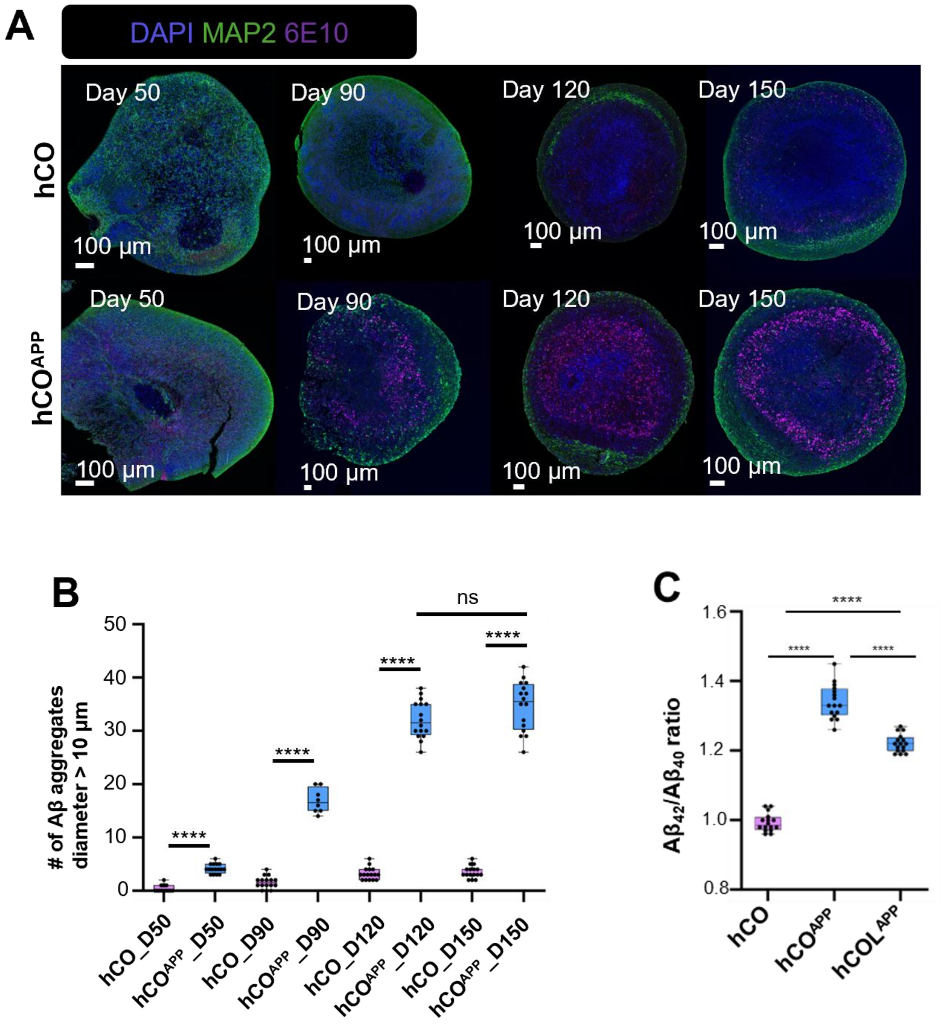


**Figure S4. Accumulation of Aβ pathology in hCO$^{APP}$.** (**A**) Immunostaining for Aβ deposition (6E10 antibody staining) and MAP2 neurons in hCO and hCO$^{APP}$ at developmental days 50, 90, 120, and 150. (**B**) Quantification of Aβ aggregates in organoid variants indicating a significantly higher number of Aβ aggregates with a diameter exceeding 10 µm withing hCO$^{APP}$ compared with control hCOs at all stages. Box plots show the median with interquartile range (n=16 organoids per group; ns=not significant, p=0.0903; ****: p<0.0001, Mann-Whitney U-test). (**C**) ELISA measurements of formic-acid–extracted Aβ1-42 and Aβ1-40 in day-120 organoids, expressed as the Aβ42/Aβ40 ratio in the insoluble fraction and normalized to total protein. hCO$^{APP}$ organoids show a higher Aβ42/Aβ40 ratio compared with control hCOs (n=16 organoids per group, confirming increased aggregation-prone Aβ species in the hCO$^{APP}$ model. ****: p<0.0001; Mann–Whitney U-test.

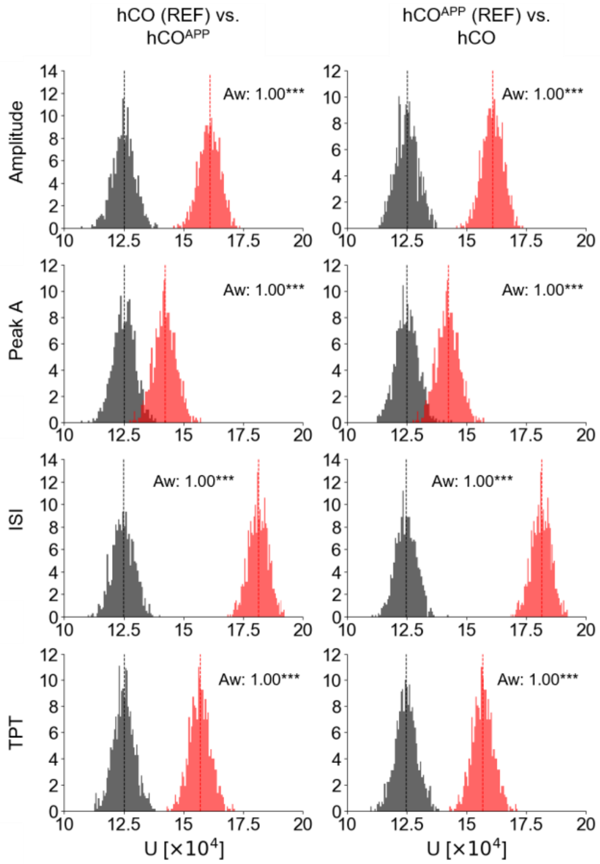


**Figure S5. Intra- and inter-class U-statistic distributions for hCO and hCO$^{APP}$ with reference class reversal.** Each row corresponds to a spike feature. The left column shows comparisons with hCO as the reference class (REF), and the right column shows comparisons with hCO$^{APP}$ as the reference. Gray histograms depict intra-class U-statistic distributions for the REF, while red histograms depict inter-class U-statistic distributions between the REF and referred class. Each U-statistic value is computed from two randomly sampled groups of 500 spikes drawn either from the same class (intra-class) or from different classes (inter-class). Each histogram is based on 1,000 repetitions, yielding 1,000 U-statistic values. *** indicate $p < 0.001$ (two-tailed Mann–Whitney U test).

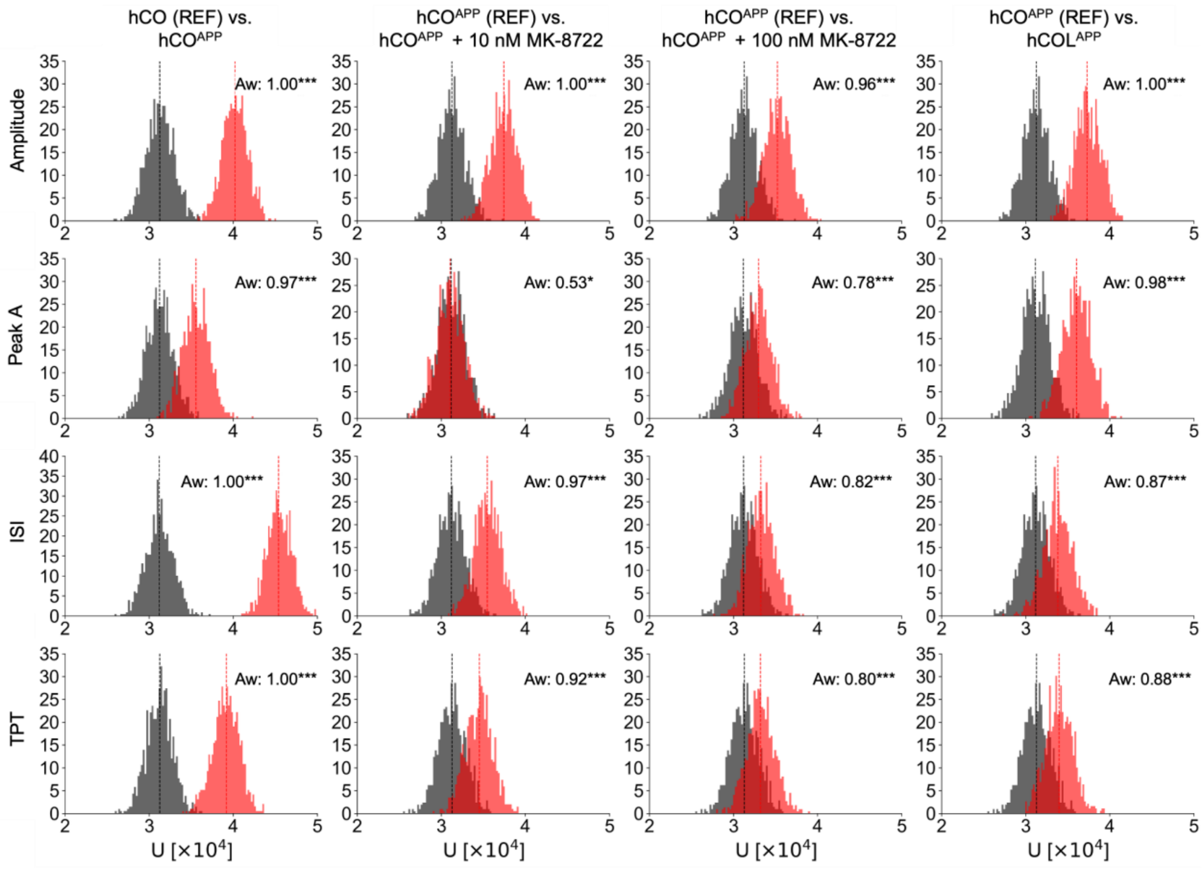


**Figure S6. The distributions of intra- and inter-organoid features using 250-spike blocks.** Each row corresponds to one spike feature, and every column corresponds to a different pair of classes. Gray histograms show the intra-class U-statistic distributions for the reference class (REF), and the red histograms show the inter-class U-statistic distributions, between the reference and the referred classes. Each U-statistic value compares two randomly sampled 250 spikes from the same class (intra-class distribution) or from two different classes (inter-class distribution), and every histogram is based on 1,000 repetitions yielding 1,000 U-statistic values. */***: p<0.05/0.001, two-tailed Mann-Whitney U-test.

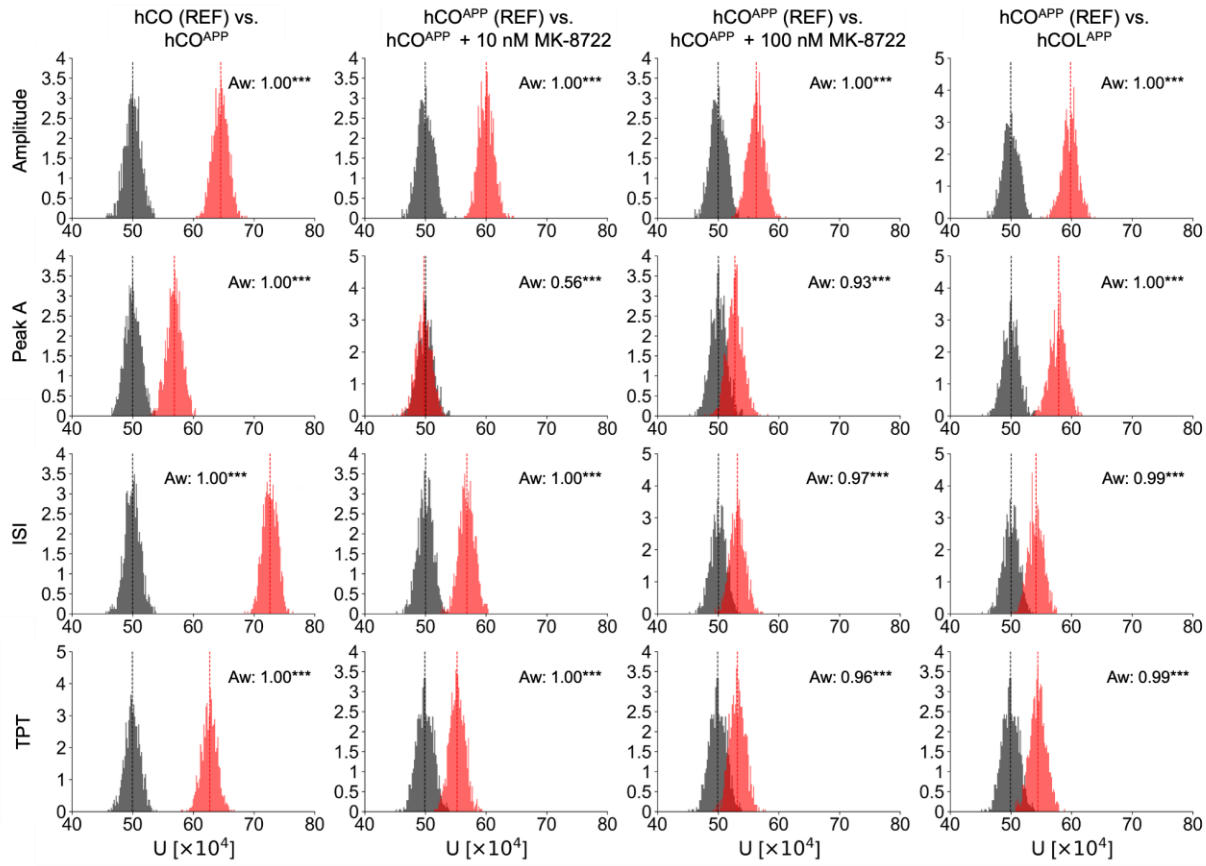


**Figure S7. The distributions of intra- and inter-organoid features using 1,000-spike blocks.** Each row corresponds to one spike feature, and every column corresponds to a different pair of classes. Gray histograms show the intra-class U-statistic distributions for the reference class (REF), and the red histograms show the inter-class U-statistic distributions, between the reference and the referred classes. Each U-statistic value compares two randomly sampled 1,000 spikes from the same class (intra-class distribution) or from two different classes (inter-class distribution), and every histogram is based on 1,000 repetitions yielding 1,000 U-statistic values. ***: $p<0.001$, two-tailed Mann-Whitney U-test.

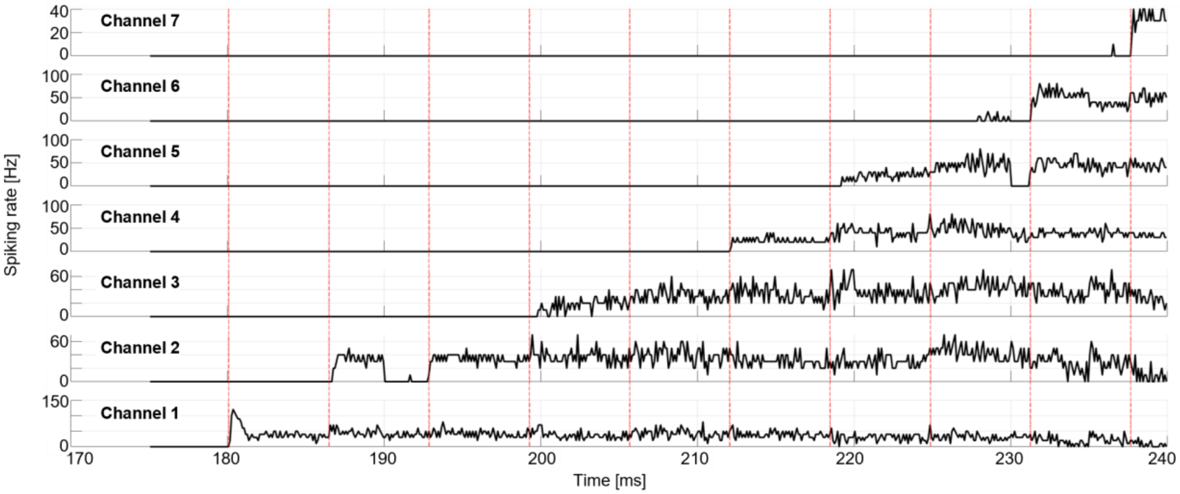


**Figure S8. Spike rate changes during probe insertion.** Spike rate was computed for each of seven adjacent channels on a single probe shank during automated insertion. The channels are ordered by the physical position along the shank, with Channel 1 closest to the probe tip, penetrating the organoid first. The motor was programmatically advanced in 20 µm increments. The onset of each movement is indicated by a vertical red line.

| | | hCO_D50 n = 10 | hCO_D115 n = 6 |
|---|---|---|---|
| Filtering (#-post-filtering) | Loaded | 20569 | 26395 |
| | UMI filtered | 16455 | 21116 |
| | Mitochondrial filtered | 15501 | 19721 |
| | Ribosomal filtered | 15345 | 19523 |
| | Gene complexity filtered | 14835 | 18821 |
| | Singlets (Analyzed) | 13226 | 17020 |

**Table S1**. Number of cells used in scRNAseq for hCO at day 50 and day 115.

| Test session | Duration [s] | Spikes | Blocks | True label | Weighted block prediction ($\hat{y}$) | Session prediction |
|---|---|---|---|---|---|---|
| hCO-6 | 3752 | 6,974 | 13 | 0 | 0.020 | 0 |
| hCO-3 | 3444 | 7,281 | 14 | 0 | 0.162 | 0 |
| hCO-15 | 3031 | 43,036 | 86 | 0 | 0.173 | 0 |
| hCO-7 | 3392 | 91,789 | 183 | 0 | 0.415 | 0 |
| $hCO^{APP}$-23 | 8955 | 1,692,234 | 250 | 1 | 0.982 | 1 |
| $hCO^{APP}$-13 | 3552 | 436,978 | 250 | 1 | 0.982 | 1 |
| $hCO^{APP}$-19 | 3367 | 896,408 | 250 | 1 | 0.979 | 1 |
| $hCO^{APP}$-5 | 4865 | 88,266 | 176 | 1 | 0.909 | 1 |
| $hCO^{APP}$-6 | 3436 | 23,794 | 47 | 1 | 0.741 | 1 |
| $hCO^{APP}$-3 | 4423 | 167,319 | 250 | 1 | 0.709 | 1 |
| $hCO^{APP}$-1 | 3624 | 19,697 | 39 | 1 | 0.623 | 1 |

**Table S2. Session-level scores for hCO and $hCO^{APP}$ test sessions yield perfect classification.** Sessions are ordered by decreasing confidence within each class group (same order as **Fig. 5D**), based on the weighted average of block-level predictions. The optimal threshold was determined from training/validation data as 0.416. The duration and number of spikes pertain to the full session, whereas the number of blocks is decimated to 250 for longer sessions.
